# From the outer ear to the nerve: A complete computer model of the peripheral auditory system

**DOI:** 10.1101/2022.04.23.489223

**Authors:** Ondrej Tichacek, Pavel Mistrík, Pavel Jungwirth

## Abstract

Computer models of the individual components of the peripheral auditory system – the outer, middle, and inner ears and the auditory nerve – have been developed in the past, with varying level of detail, breadth, and faithfulness of the underlying parameters. Building on previous work, we advance the modeling of the ear by presenting a complete, physiologically justified, bottom-up computer model based on up-to-date experimental data that integrates all of these parts together seamlessly. The detailed bottom-up design of the present model allows for the investigation of partial hearing mechanisms and their defects, including genetic, molecular, and microscopic factors. Also, thanks to the completeness of the model, one can study microscopic effects in the context of their implications on hearing as a whole, enabling the correlation with neural recordings and non-invasive psychoacoustic methods. Such a model is instrumental for advancing quantitative understanding of the mechanism of hearing, for investigating various forms of deafness, as well as for devising next generation hearing aids and cochlear implants.

## 1 Introduction

Proper understanding of the mechanism of hearing and especially the cochlear function is important for the advancement of the compensation of hearing loss, including the design and setup of hearing aids and cochlear implants. While some psychophysical properties of the human cochlea have been thoroughly studied and a sufficient amount of data is available (Robles and Ruggero 2001), others are hardly accessible due to the invasive nature of the measurements. In these cases, data may be obtained, at least to a certain extent, from animal models. Importantly, computer modeling can provide insight to particular problems, where experiment is not feasible, and help to establish a detailed quantitative understanding of the system as a whole.

The mammalian peripheral auditory system consists of the outer, middle, and inner ears (the latter including the cochlea) and the auditory nerve leading to the cochlear nucleus, where in turn the central auditory nervous system (CNS) starts. Its main function is to detect the sound and convert it to neural activity to be processed by the CNS. Considering the immense range of sound frequencies of 20 Hz up to 20 kHz and intensities from 0 to ∼100 dB sound pressure level (SPL), the function of the human auditory periphery is enabled by highly specialized multi-cellular and intracellular structures, often unique to the auditory system, that are responsible for key mechanisms such as mechanosensitivity, amplification, and adaptation. Besides that, there are many other processes on the peripheral auditory pathway that enable the mechanism of hearing (see Fig. 1), all of which need to be included in the computer model in order to obtain an accurate description of the physiological reality.

**Figure 1:**
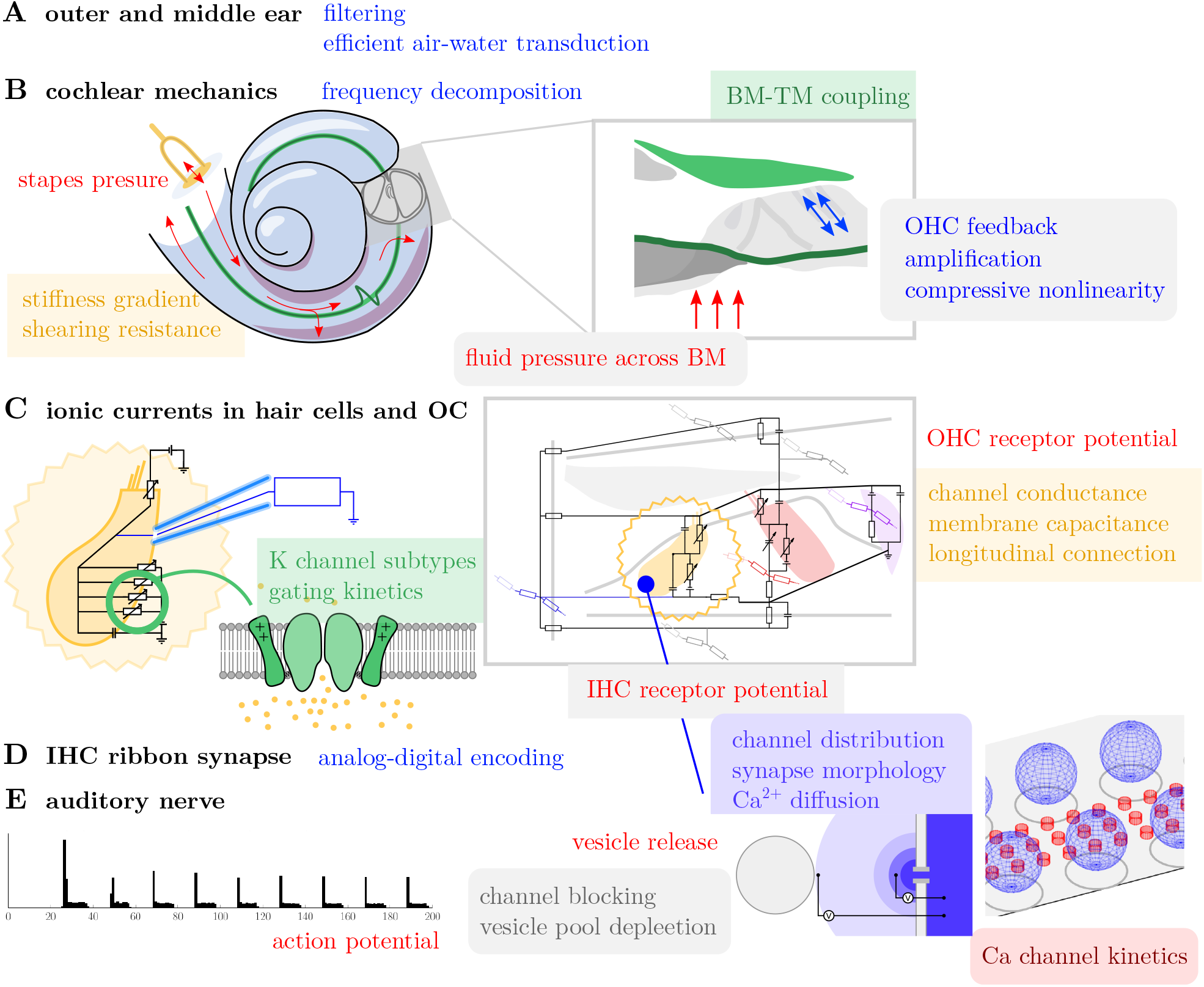
Model blocks and the mechanism of hearing. **A**: Acoustic waves pass through the outer ear and middle ear, which effectively conveys the vibration from air to the cochlear fluids. **B**: Pressure oscillations in the cochlear fluids are spatially decomposed across the basilar membrane (BM), coupled to the tectorial membrane (TM) depending on their frequency in the first step to sound encoding. **C**: Motion of the organ of Corti (OC) and subsequent deflection of the stereocilia in the sensory cells opens ion channels and induces transduction of ionic current. **D**: Change of the receptor potential in the inner hair cells (IHC) triggers release of synaptic vesicles and thus evoked action potential in the afferent nerve fibers finalizes the passage from analog to “digital” signal (**E**). In the outer hair cells (OHC), the receptor potential causes conformation change of a voltage-sensitive protein prestin which translates to macroscopic contraction and elongation of the cell and a consequent force acting on the organ of Corti. When in phase, this force can act as an amplifier of the cochlear mechanics, being responsible for a sharper tuning and increased level detection range through the compressive nonlinearity.

Computer modelling of the auditory periphery has been a busy field in the past decades, see for instance Refs. (Bruce 2006; Bruce et al. 2018; Heil et al. 2011; Mammano and Nobili 1993; Meddis 1986; Mistrík, Mullaley, et al. 2008; Pascal et al. 1998; Strelioff 1973; Sumner et al. 2002; Verhulst, Altoè, et al. 2018; Verhulst, Dau, et al. 2012; Vetešník and Gummer 2012) and Ref. (Vecchi et al. 2022) for a recent comparison of existing models. Most of these models focus on individual parts of the auditory periphery and in several instances need to be updated to reflect the most recent experimental findings both on humans and animal models. The level of description varies greatly among published models. For example, models of cochlear mechanics range form filter-bank models to detailed three-dimensional models including cochlear micro-mechanics. All of these models have their place in the auditory research and are generally used for different purposes. Some can speedily mimic various experimental recordings (e.g., the neural activity in response to a sound stimulus), while other provide a deeper insight into mechanisms underlying important aspects of hearing, such as hair cell function or the cochlear amplifier. Models combining the breadth of simpler models with a detailed microscopic description of complex models are largely missing, with only a few exceptions Verhulst, Altoè, et al. (2018). Such complete, detailed models are important, for example, for assessing or predicting the implications of modifications in partial hearing mechanism on hearing in general. Here, we have moved forward in two ways: First, by creating a framework of models, each of which being more advanced than its predecessor in the literature and, second, by organically connecting these models into a single system. Tackling computationally expensive tasks by efficient implementation has allowed us to avoid compromises concerning the level of description of the individual components of the peripheral auditory system.

In this paper, we thus present a physiologically well-justified computational model, with its core being a complete cochlear partitioning with a detailed description of electrical excitability of individual hair cells. We couple it with models of outer and middle ears, a model of the cochlear mechanics, a conceptually novel model of the ribbon synapse of the inner hair cell, and with a model of the auditory nerve. The present complete model of the peripheral auditory system is constructed in a “bottom-up” way, based on the appropriate laws of physics and physiologically relevant measurable properties of its building blocks, avoiding thus “black-box” heuristic approaches. Where human data are not available, animal data have been used instead, coming exclusively from mammalian species such as the guinea pig. The model as a whole has been tested for a wide range of input frequencies (up to 8 kHz) and sound levels (up to 100 dB SPL), reproducing well experimental data whenever available and otherwise yielding results consistent with existing models.

## 2 Theory and Results

From the outer ear to the nerve, the sound waves are conveyed, processed, and encoded by several mutually interconnected and cooperating biological systems. The present model is designed in the same way – as an aggregate of coupled models of the outer ear, middle ear, the cochlear mechanics, the ionic currents within the cochlea and the hair cells, the ribbon synapse, and the auditory nerve. In the following sections, key novel aspects of the individual models and their interconnections are described, while an exhaustive description is provided in the Methods section and in the supplementary information. Numerical values of model parameters are available within the source code, which may be obtained upon request.

While in this paper we focus primarily on the methodology, we also provide a proof of concept of application of the model for the following examples: (1) we investigate the nonlinear effects of potassium channel kinetics coupled with nonlinear OHC capacitance using our model which includes the tonotopic specialization of hair cells, (2) we model a partial dysfunction of IHC and OHC stereocilia, (3) we study the coupling of the cochlear amplifier and the OHC receptor potential, (4) we reproduce experimental threshold shifts (Chen et al. 2020) using a model with a damaged cochlear amplifier, and (5) we simulate maturation of the ribbon synapse of the IHC, offering a microscopic quantitative insight and verifying the conclusions from experiments (P. F. Vincent et al. 2018).

### 2.1 Acoustic transduction through outer and middle ears

The outer ear transfer characteristics are implemented in the present model via a set of independent band-pass filters as proposed by Meddis (2011) and calibrated to experimental data of Mehrgardt and Mellert (1977), see Fig. 2A. Following Pascal et al. (1998) the middle ear is represented as a lumped-element model of selected structures — namely the eardrum, the middle ear cavities, malleus, incus, stapes, and the cochlea. Here, we recalibrated the model to recent experimental tympanic membrane pressure-stapes velocity data by Chien et al. (2009), as demonstrated in Fig. 2B.

**Figure 2:**
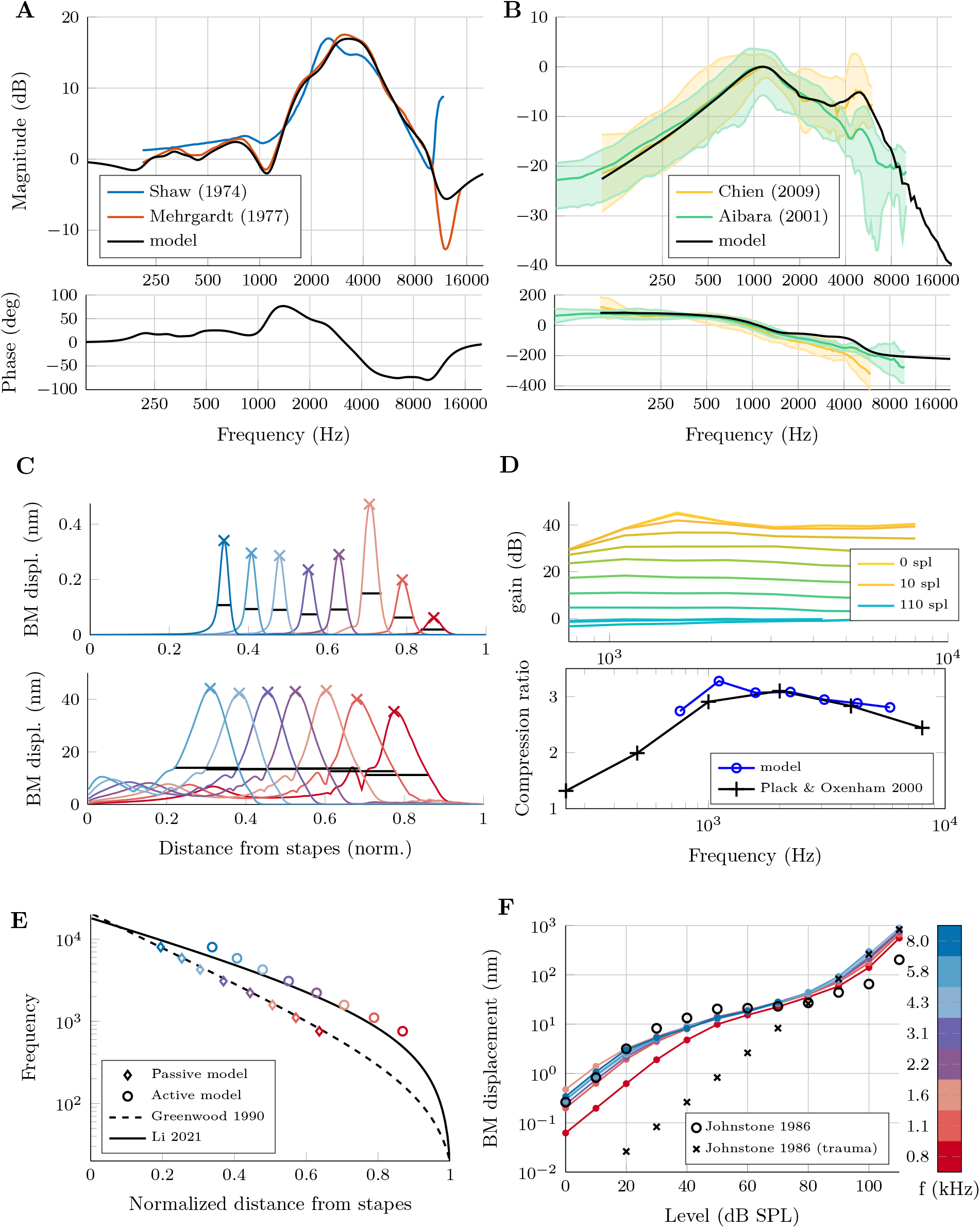
Mechanics of the outer and middle ears, and the cochlea: Comparison between simulated data and experiments on humans or animal models. **A**: Pressure-pressure transfer function of the outer ear model compared to experimental data by Shaw (1974), and Mehrgardt and Mellert (1977). Experimental phase shifts are not available. **B**: Pressure-velocity transfer function of the middle ear model with experimental data by Chien et al. (2009) and Aibara et al. (2001). **C-F**: Basilar membrane response as calculated by the model; **C**: Traveling wave envelopes for of BM oscillations for pure tones of 0.8 to 8 kHz at 0 dB SPL (top) and 80 dB SPL (bottom). **D**: Measured relative gain of the active vs. passive model (top) and the mean compression ratio between 50 an 80 dB SPL (bottom) compared to psychoacoustic experimental data by Plack and Oxenham (2000). **E**: characteristic frequency (CF) as a function of the position along the BM as reproduced by the model (in both active and passive modes) compared to experimentally based CF curves by Greenwood (1990) and Li et al. (2021). **F**: Input-output curves of BM displacement vs. sound level for range of frequencies from 0.8 to 8 kHz compared to experimental data by Johnstone et al. (1986).

The model can closely reproduce the experimental data up to ∼10 kHz (outer ear) and ∼8 kHz (middle ear). For higher frequencies, individual experimental datasets are mutually less consistent or even missing, which complicates accurate validation of the model in the very high frequency range.

### 2.2 Cochlear mechanics

The first step of sound encoding occurs as the basilar membrane spatially decomposes the sound wave based on its frequency components. This is enabled primarily by the gradual change in its stiffness and other mechanical properties along the tonotopy axis, i.e., from the base to the apex of the basilar membrane.

Here we employ a nonlinear 2-dimensional model of cochlear mechanics based on that by Vetešník and Gummer (2012). A major step forward in the present work is an advanced description of the cochlear amplifier employing a direct feedback from the subsequent model of the electric currents within the cochlea (see below). The amplification with respect to a passive model can reach (depending on the stimulus frequency) ∼45 dB SPL (Fig. 2D), while maintaining the zero crossing condition (Supplementary Fig. S-1). The compression ratio of the input/output curves of the BM displacement (Fig. 2F) is comparable with psychoacoustic experimental data (Plack and Oxenham 2000). We have also reparametrized the model to reproduce the human tonotopic maps Greenwood (1990) and Li et al. (2021) (Fig. 2E), which is crucial for correct assessment of tonotopy specialization of hair cells.

### 2.3 Cochlea as a large-scale electrical circuit

In this section, we provide the description of the cochlear model in terms of a large-scale electrical circuit. In particular, we summarize the updates to the circuit design compared to earlier models in the literature, focusing on the present advanced description of the hair cells. These improvements concern primarily the implementation of (i) the conductance of the stereocilia through the mechanosensitive channels, (ii) the voltage-dependent conductance of the basolateral membranes dominated by the potassium channels, (iii) potassium channel sub-types with different gating kinetics, and (iv) the non-linear part of the capacitance of the cell membranes of the outer hair cell (OHC).

The modelling approach adopted here follows a general concept of a lumped-element model, first introduced by Strelioff (1973) for the analysis of the intracochlear distribution of electrical potentials. Such approach was used later also for modelling of a cochlea implanted with a multielectrode stimulating array (Suesserman and Spelman 1993) and of a three-dimensional current flow and its effect on the amplification of sound (Mistrík, Mullaley, et al. 2008), as well as for modelling of reduced OHC electromotility associated with connexin-related forms of deafness (Mistrík and Ashmore 2010).

In the present model, the cochlea is approximated by interconnected electrical circuits representing the radial flow of an electric current at each cross-section, as well as the longitudinal current flow between cross-sections. Compared to previous models (Mistrík and Ashmore 2010), the number of longitudinally connected cross-sections was increased from 300 to 3000, corresponding to a typical number of inner hair cells (IHCs) in the human cochlea (the number of OHCs being about four times larger). In this way, each cross-section accounts for the radial current flow through each row of hair cells in the human organ of Corti. For specific situations (e.g. large-scale simulations that would be limited by the amount of produced data) the number of cross-section can be adjusted and the circuit automatically coarse-grained. An equivalent electrical circuit of a single cross-section of the cochlear partition is presented in Figure 3A. Given experimentally accessible input parameters, such as the characteristic conductances and capacitances of the cells of interest and the electrical/electrochemical potentials, the model provides currents and transmembrane potentials at each part of the system (Supplementary Fig. S-3). Its principal advantages are revealed upon coupling with a model of cochlear mechanics that simulates the motion of the basilar membrane and the stereocilia deflection. Namely, these motions can be directly translated into conductance changes of the hair cell stereocilia and, consequently, into evoked currents and potentials throughout the system. In this way, receptor potentials in the hair cells, generated in response to arbitrary sound stimuli, can be realistically estimated (Figures 3B-C, 5D, Supplementary Figures S-2, S-3, and S-4).

**Figure 3:**
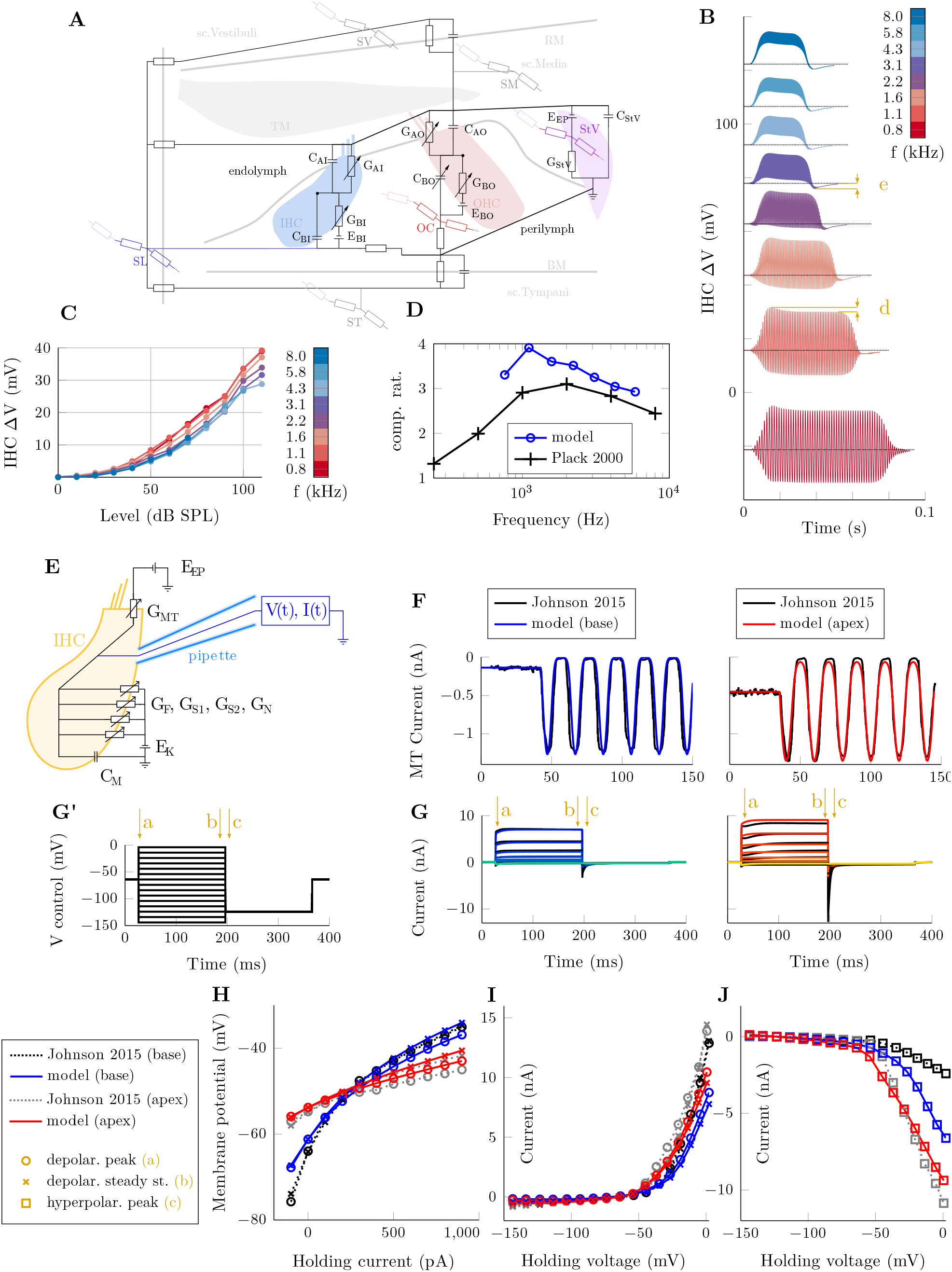
Ionic currents within the cochlea: **A**: An equivalent electrical circuit of a cochlear cross section. Electrical properties of the apical and basolateral parts of inner and outer hair cell membranes are described by the corresponding resistors (conductance *G*) and capacitors (*C*). The reversal potential arising from the difference between intracellular and extracellular potassium concentrations is represented by a voltage source (battery). Each IHC is accounted for individually, while four OHCs in a cross section are represented by one OHC with scaled values of their *G* and *C* elements. Stria vascularis (StV) is represented by a corresponding resistor and capacitor, and the presence of the endocochlear potential at scala media (SM) by a battery. Other cellular structures in the organ of Corti and spiral limbus (SL) are represented by resistors. The Reisner’s and basilar membranes separating the scala vestibuli (SV), media, and tympani (ST) are represented by a resistor and a capacitor. Each of the three scalae, the spiral limbus, organ of Corti (OC) and the stria vascularis are longitudinally connected to other cross sections via resistors. Some of the elements are variable; the resistors *G*_AI_ and *G*_AO_ are modulated by the stereocilia deflection, *G*_BI_ and *G*_BO_ are regulated by voltage and the kinetics of the ion channels they represent, and the value of *C*_BO_ includes the nonlinear capacitance, which is voltage dependent. Moreover, the hair cell basolateral resistors are composed of four independent resistors representing different subtypes of expressed potassium channels, see panel E. **B-D**: IHC receptor potential (∆*V*) as computed by the full 3D-circuit model in response to pure tones of varying frequency and level: **B**: Waveforms of IHC ∆*V* at the maximal magnitude cross-section at 70 dB SPL exhibit expected properties such as AC/DC ratio decrease with stimulus frequency. The peak value occurring shortly after the stimulus onset is larger than the steady-state maximum (indicated by “d” in the figure) and similarly, the potential short after the stimulus end is hyperpolarized compared to the resting state (indicated by “e”). Both of these effects are caused by the non-instantaneous response of the channels to the change of the membrane potential (i.e., the channel kinetics). See supplementary figure S-4 for more details on the effect of the stimulus onset. **C**: Input-Output curves of the peak IHC receptor potential exhibit a compressive nonlinearity similarly to the BM displacement in the 40 to 80 dB SPL range. Another compression occurs for high sound levels due to the displacement-induced conductance saturation of the mechanosensitive ion channels in the stereocilia tips. Consequently, as can be seen from the figure, the receptor potential is almost linear with respect to the the sound level in dB SPL from about 40 dB. The receptor potential is diminished for higher frequencies due to a low pass filtering of the RC circuit (i.e., the membrane plus channel system), see also panel C. **D**: The mean compression ratio between 50 an 80 dB SPL compared to psychoacoustic experimental data by Plack and Oxenham (2000), is increased compared to the BM displacement (Fig. 2). **E-J**: Model of an isolated hair cell. **E**: Diagram of the model in the whole-cell patch-clamp configuration. **F**: The mechano-electrical transduction (MET) current elicited by stereocilia displacement in the isolated cell model configuration compared to experimental data measured by Johnson (2015) with stereocilia displaced using a fluid jet. **G**: Example of a voltage-clamp protocol (depolarizing and hyperpolarizing voltage steps of different magnitude (G’)), simulation and experiment by Johnson (2015). The arrows **a, b, c** denote features of the response to the current clamp (**H)** and voltage clamp (**I, J**) protocols used by (Johnson 2015) to differentiate between basal and apical cells. **E**: Peak (a) and steady state (b) voltage in response to a step in control current. **F**: Peak (a) and steady state (b) current in response to voltage step. **G**: Amplitude of tail currents (c) in response to a hyperpolarizing step.

### 2.4 Electric currents in a single hair cell

Parametrization of the electrical properties was adopted from Strelioff (1973) regarding the conductivity of fluids in the scala vestibuli (SV), scala media (SM), scala tympani (ST), and spiral limbus (SL) and from Mistrík, Mullaley, et al. (2008) concerning the organ of Corti (OC), being based on resistive and capacitive properties of the cellular constituents. The numerical values of electric elements were updated to reflect the latest experimental data (Johnson 2015; Johnson, Beurg, et al. 2011), particularly for the OHC and IHC specialization along the tonotopic axis. The calibration was aided by a supplementary model of an isolated hair cell, reproducing in-vitro electrophysiological experiments (Fig. 3E-J). Such an approach is necessary, as experiments with isolated cells cannot be correctly reproduced solely by a model of the whole OC, and similarly, in-vivo characteristics of hair cells (e.g. resting or receptor potentials) are affected by electrical properties of other sub-structures of OC and neighboring cross-sections of the cochlear partiton and thus cannot be reliably estimated with models of isolated hair cells. To illustrate these effects, we calculated the IHC receptor potentials using the model of the full cochlear partition and compared it with a model of an isolated IHC, similar to the one reported by Dierich et al. (2020). Both models used identical values of electrical elements and we also included an artificial endocochlear potential in the model of an isolated IHC and enforced the same deflection of stereocilia and other simulation conditions. We see significant differences between the results of the two models, in particular, a shift in the resting potential (on average) by 2.5 mV and a difference in the amplitude of the receptor potential by up to 0.2, 6.1, or 21.0 mV (i.e., by 100–300 %) when stimulated by a 2 kHz pure tone of 0, 40, or 80 dB SPL. We also performed simulations for other frequencies between 1 and 8 kHz with qualitatively identical results.

### 2.5 Nonlinear effects – potassium kinetics and OHC capacitance

The non-linearity and voltage-dependence of membrane resistances due to the presence of voltage-gated ion channels was implemented together with the potassium channel gating kinetics that accounts for a non-instantaneous response of the population of the channels. Channel kinetics were modeled similarly as in Lopez-Poveda and Eustaquio-Martín (2006), but with the model being extended to include different potassium channel subtypes (K_f_, K_n_, and multiple variants of K_s_, their distributions varying along the tonotopy axis) and calibrated to match the recent experimental data (Johnson 2015), see Fig. 3F-J. Since the gain of the cochlear amplifier driven by the cochlear electromotor prestin is in our model influenced to a large extent by the value of OHC capacitance, it is critical to determine this capacitance accurately. Therefore, the nonlinear capacitance model of the OHC basolateral membrane (Dallos and Fakler 2002; Santos-Sacchi and Tan 2020) was also implemented (see Methods). While it turns out that the changes in capacitance due to the sound-evoked changes in the receptor potential do not have much effect on the amplification, shift of the capacitance due to a shift of the steady-state (average) potential of OHC can have significant impact on the amplification. Since the steady state potential of the hair cells is not a model parameter, but is instead calculated on the fly by solving the electrical circuit, the dependence of the capacitance on the transmembrane voltage is an important factor included in the model.

The present methodology is important for properly capturing the nonlinear properties of the system. As an illustrative example, we have investigated the effect of voltage-dependent channel gating kinetics of basolateral IHC ion channels, in comparison to the commonly used linear approximation Mistrík, Mullaley, et al. (2008). Considering the voltage dependence of the potassium channel open-probability, the IHC membrane time constant is decreased upon depolarization of the cell and increased upon hyper-polarization. The non-instantaneous channel gating kinetics further modulates this effect. The affected filtering properties of the membrane–channel system can be summarized by the magnitude of the AC/DC components of the receptor potential. While for near-threshold low-frequency stimuli the present methodology and linear approximation give comparable results, with increased sound level or frequency our model shows that the magnitude of the AC component is significantly underestimated by the linear approximation (Supplementary Fig. S-6). Note that for AC/DC analysis to be indicative of filtering effects of the system the DC component must naturally be present. Since the DC component of the receptor potential is the product of the MET channel gating nonlinearity, which gains on significance with SPL, it only appears for signals far from threshold (the DC component is approximately 1, 10, and 60 % at 5, 35, and 65 dB SPL, 8 kHz tone, see Fig. S-6-A). However, for near-threshold stimuli, the voltage-independent basolateral channel gating kinetics is a good approximation, thus no effects on the filtering properties of the system are to be expected.

Another example of the effects of the non-instantaneous hair cell channel kinetics concerns the stimulus envelope. Namely, the slope of the onset affects the later response – a short (steep) onset causes temporarily higher depolarization magnitude than a long onset. A short vs. long offset of a stimulus has a similar effect; see Fig. 3B and Supplementary Fig. S-4. The effect mimics adaptation of the neural response caused by stockpiling/depletion of the readily-releasable vesicles by the ribbon synapse. Since depolarization eventually triggers synaptic release events while hyperpolarization attenuates them, the hair cell channel kinetics can be partially responsible for the observed adaptation (its short-term component).

### 2.6 Mechano-electrical transduction

Deflection of the IHC and OHC stereocilia opens MET channels and increases the instantaneous conductance of the hair cells. Analogously, in the present model, the electrical circuit is modulated by the motion of the basilar and tectorial membranes. We model the OHC MET open probability as a two-level Boltzmann sigmoid function as in Mistrík, Mullaley, et al. (2008). In contrast to Mistrík, Mullaley, et al. (2008) where it is coupled to the BM displacement, in the present model the MET open probability is governed by explicitly calculated deflections of stereocilia, which is physiologically more realistic. We model the IHC MET open probability similarly to Verhulst, Altoè, et al. (2018) using a three-level Boltzmann sigmoid. For both OHC and IHC, the MET transduction parameters have been updated to reflect experimental data by Jia et al. (2007) and Johnson (2015), in particular to reproduce the tonotopical variations.

Dysfunction of the MET channels, which can lead to hearing loss, and can be caused by a variety of factors, including genetic mutations (Xiong et al. 2012), age-related degradation, noise exposure, ototoxic drugs, and others. When investigating MET channel dysfunction, it is advantageous to understand the expected response of the system. This information can be correlated with global hearing characteristics such as hearing threshold, especially when using non-invasive measurements.

Figure 4 shows the effect of partial damage to the inner hair cell (IHC) or outer hair cell (OHC) stereocilia, such as from noise exposure. Differences in the cochlear function degradation caused by dysfunction of MET channels in the OHC and IHC were observed. Cochlear function was approximated for simplicity by the amplitude of the IHC receptor potential. In the case of impeded response due to noise damage, the affected portion of the cochlea was significantly narrower when IHCs were damaged. For the case when OHC were affected, the response was asymmetric and shifted relative to the center of the noise band. Such differences could potentially be used for dysfunction diagnostics, possibly even by combining different factors using parameter-inference.

**Figure 4:**
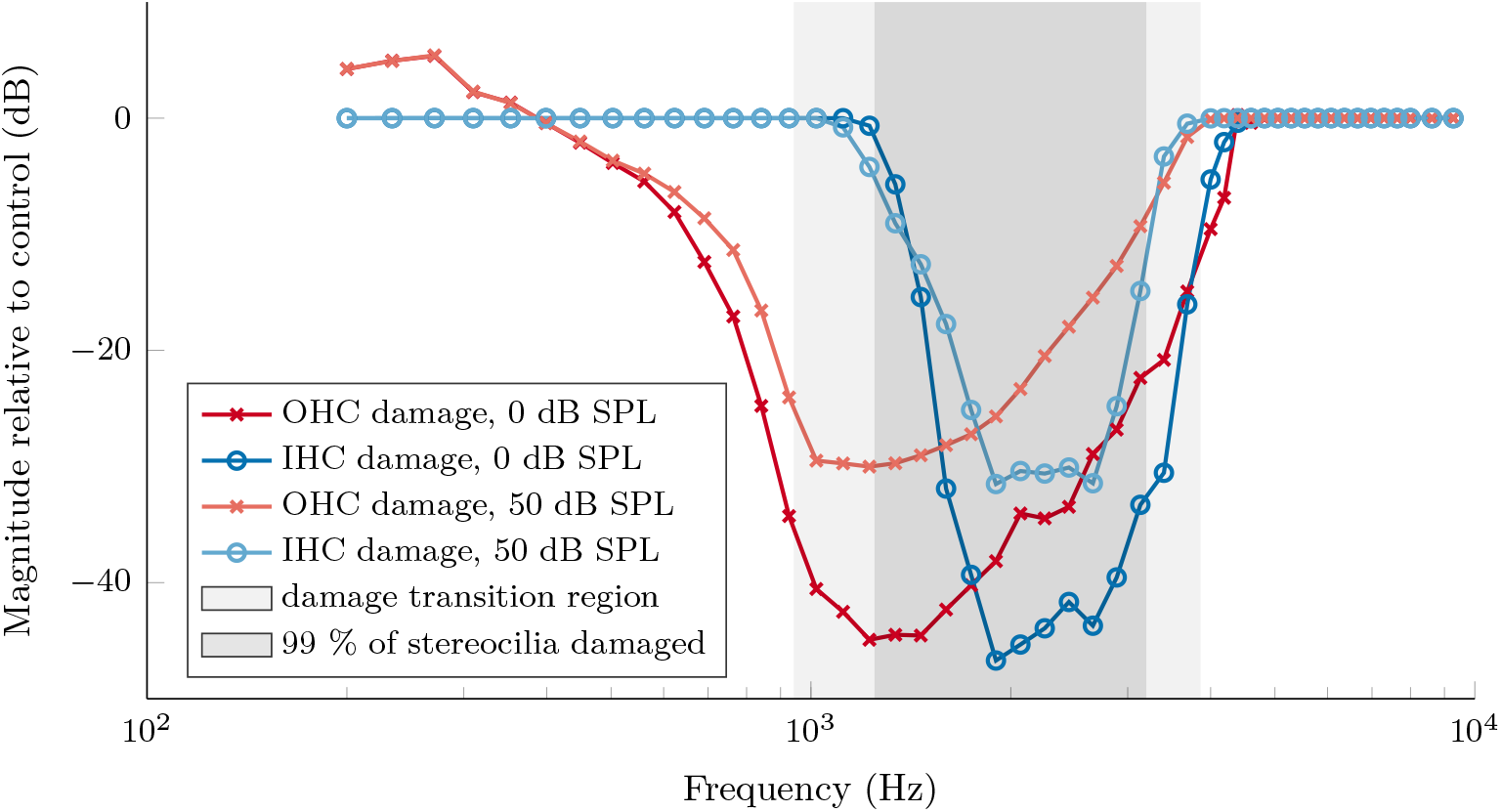
Dysfunction of IHC and OHC stereocilia: Simulating a damage to IHC and OHC stereocilia, we have decreased selectively the conductance of MET channels to 1% in the region corresponding to best frequency of 1.25 to 3.20 kHz with a smooth transition to unaffected stereocilia below 0.94 kHz and above 3.86 kHz. In the present two sets of simulations, only either IHCs or OHCs were damaged, the other unaffected to allow for comparison of the effect. We performed independent time-domain simulations using pure tones from 0.2 to 9.3 kHz at 0 and 50 dB SPL and recorded the IHC receptor potential (for both damaged-IHC and damaged-OHC simulation sets). We analyzed the magnitude of the response in the terms of the amplitude of the IHC receptor potential, serving as a proxy to the neural response, with respect to a control simulation when neither OHCs nor IHCs were damaged. The differences in the shape of the magnitude re. control (especially the asymmetry and width) can reveal the origin of hearing loss (in this example damaged stereocilia of IHCs or OHCs).

### 2.7 Cochlear amplifier

Nobili and Mammano (1996) investigated the nonlinear cochlear mechanics and introduced an “active amplifier” term in their model. This term was theoretically derived from the response of prestin to the OHC receptor potential, but the potential itself was not explicitly calculated. Instead, the active force term was made proportional to the stereocilia deflection, saturating with increasing intensity, and calibrated so that it mostly cancels the shearing resistance. The same approach was adopted also in the later model by Vetešník and Gummer (2012).

In order to make the model more realistic and to simplify it from a theoretical point of view, we replaced the indirect estimation of the OHC receptor potential by an exact computation from the electrical model of cochlear partition. The main advantage of this approach is the direct inclusion of effects produced by the present complex and fine-grained, longitudinally coupled electrical model, in particular, the voltage sensitivity and dynamics of the ion channels, tonotopic gradients in OHC conductance and capacitance, the nonlinear OHC capacitance, as well as additional filtering effects caused by other components of the model. Consequently, there are three main sources of nonlinear behaviour in the present amplifier model – the active force as a function of the receptor potential, the voltage-dependent conductance of basolateral K^+^ channels, and the MET conductance governed by the stereocilia deflection.

Increasing progressively the amplification level of the model allowed us to investigate the amplification mechanism (see Fig. 5A-C). For amplification to occur, the force has to be in phase with the BM velocity. Due to the phase roll-off of the motion of BM, this happens at a slightly different tonotopic location than what would correspond to the characteristic frequency of the passive vibration. When the BM velocity and the amplification force are in opposite phases, attenuation occurs instead of amplification, which leads to sharpening of the response (Fig. 5A vs. B and also Fig. 2C). While in the original model by Vetešník and Gummer (2012) the amplification force is by definition in phase with the stereocilia deflection, in the present model the membrane capacitance together with the kinetics of the basolateral K^+^ channels induce a frequency and amplitude-dependent phase lag. Consequently, the tonotopic map of the active model is shifted towards the apex compared both to a passive model and to a model with implicitly calculated OHC potential, see Fig. 2C,E and 5B vs. C, which can explain the shift in experimental tonotopic maps (Greenwood 1990; Li et al. 2021).

**Figure 5:**
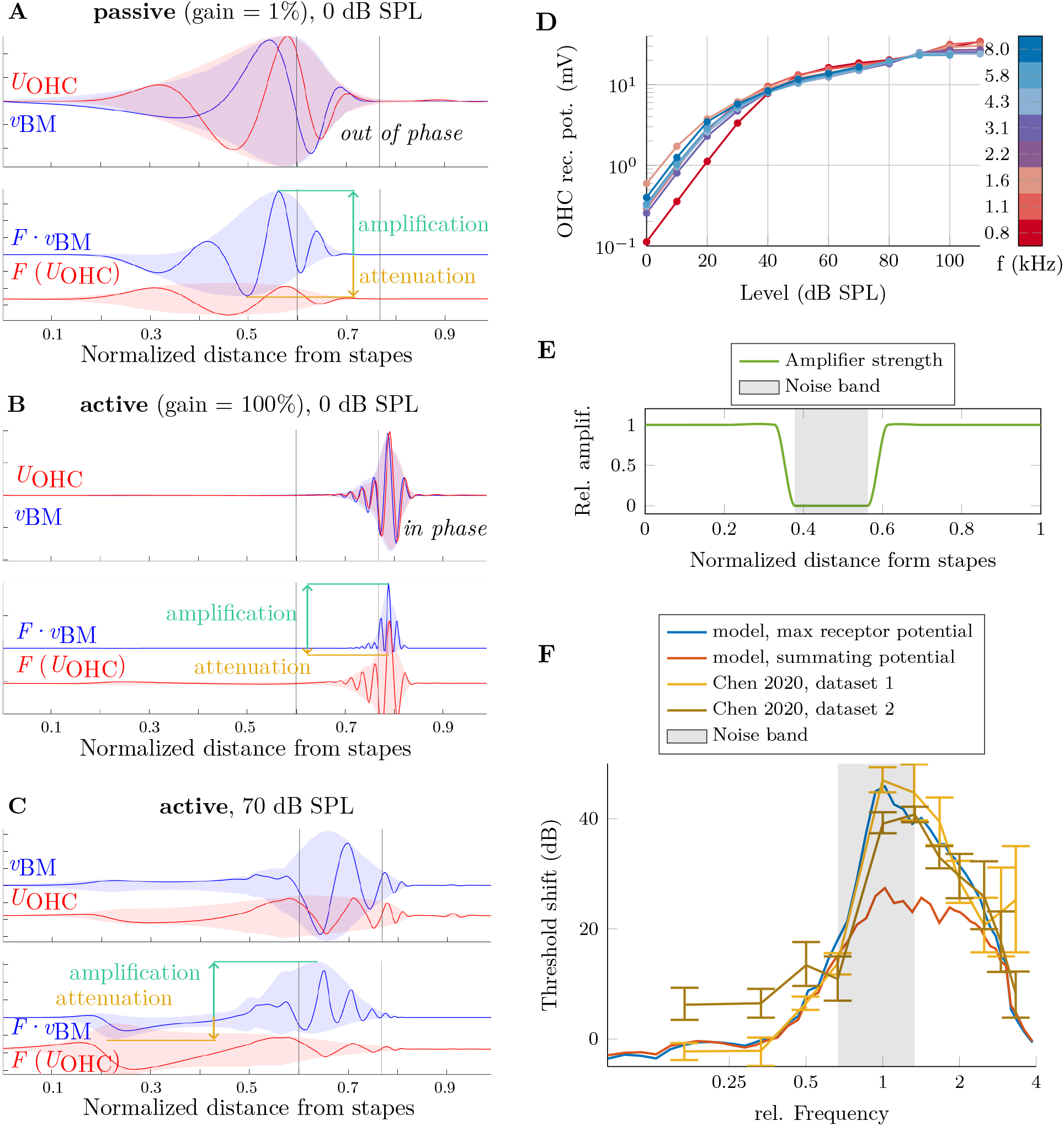
The mechanism and effect of the cochlear amplifier. **A-C**: The OHC receptor potential *U*_OHC_, the BM velocity *v*_BM_, the amplification force *F* and the amplification measure, calculated as *F · v*_BM_, all along the normalized distance from the stapes. The curves show a value at a single time point, with colored areas showing the min-max envelope throughout the simulation. All three panels show a simulation of pure tone of 1 kHz, at 0 dB SPL for (A) passive and (B) active cochlea and at 70 dB SPL for an active cochlea (C). With increasing gain, the characteristic position for a given frequency moves towards the apex, where the BM velocity and the amplifier force are correctly in phase for amplification to occur. **D**: The magnitude of the OHC receptor potential, crucial for proper function of the cochlear amplifier, is not dramatically reduced for high frequencies, as might be expected due to the low-pass filtering properties of OHC cell membrane. Instead OHC specialization along the tonotopy axis largely cancels the filtering effect, rendering the OHC receptor potential magnitude similar across frequencies. **E-F**: Modelling noise induced hearing loss – effect of the cochlear amplifier: We have exposed a model cochlea to a 1.6–3.2 kHz single octave 95 dB SPL band-noise, measuring the extent of BM oscillations for band-edge frequencies, thus determining the damage extent along the tonotopy axis, resulting in a locally reduced function of the cochlear amplifier (E). Then we performed simulations of 0.2–9.3 kHz pure tones in both normal and damaged cochlei, recording the IHC receptor potential. The figure (F) shows a comparison of the peak IHC receptor potential and estimated summating potential “damaged-to-control shift” to the experimental CAP threshold shift in mice by Chen et al. (2020) normalized to the central frequency of the noise band.

The present model has the potential to be used as a tool to correlate the experimentally accessible global hearing measures, such as the hearing threshold, with the partial effect of elemental hearing mechanisms and especially their disorders. To demonstrate this capability, we modelled the amplification in a cochlea with locally impaired amplification ability. Aiming to reproduce experimental measurements of Chen et al. (2020), who measured compound action potentials (CAP) in mice after long-term exposure to high-level octave band noises leading to severe noise-induced hearing loss, we selectively turned off the OHC amplifier and measured the IHC response in terms of the receptor potential and the summating potential. It turns out, that the experimental CAP threshold shift can largely be explained in our model by damage to the OHC only (IHCs being unaffected), including the spread of threshold shift outside the noise band; see Fig. 5E-F.

### 2.8 Response to soft sounds

Sounds near the hearing threshold inflict a peak displacement of the BM in the sub-nanometer level and the consequential MET probability change is very small correspondingly (0.7 % for 1 nm displacement). Considering the number of MET channels in a hair cell (∼300 based on the single channel conductance, Beurg et al. (2021) and Géléoc et al. (1997)), using the deterministic approach traditionally employed in models of hair cells (Dierich et al. 2020; Lopez-Poveda and Eustaquio-Martín 2006; Mistrík and Ashmore 2010), soft sounds would evoke an opening of less than one channel. In reality, the channels open stochastically and only their mean open count can be considered continuous. While the approximation coming from the continuous representation of the conductance is negligible for high-level stimuli, it can play a significant role for low-level ones. We demonstrate here that in such cases in reality at certain random time instances the IHC depolarize significantly more than in a classical deterministic model, while at other instances it does not depolarize at all. On average, however, the mean MET open probability/conductance as well as the value of the receptor potential are identical, see the Supplementary Figure S-5.

Considering that some of the elemental processes of the IHC signal transduction (Ca_V_1.3 open probability, vesicle release calcium threshold from the Ribbon synapse, etc.) are nonlinear in nature (Beutner et al. 2001; Zampini, Johnson, Franz, Lawrence, et al. 2010), this stochastic effect may be significant for the detection of signals close to the hearing threshold. It may also be the first instance, when the level is (partially) encoded as rate, such as is the case in the auditory nerve. To test this hypothesis, we investigated whether the stochastic effect is large enough to induce significant non-linearity in calcium channel kinetics in a way that can be used for near-threshold signal detection. As the mechanism of near-threshold hearing is not yet fully understood, we consider the following two possibilities: (i) the signal is detected through an increase in the cumulative Ca_V_1.3 open probability, or (ii) the signal is detected via intra-cycle differences of the Ca_V_1.3 open probability (i.e., differences between the “positive” and “negative” halves of the cycle). We implemented this into the preprocessing of the IHC synapse model by solving again an isolated IHC circuit, while including the stochastic nature of the channels. We then compared the Ca_V_1.3 open probability using the stochastic IHC receptor potential (averaged over many simulations) to the classical deterministic potential and observed a significant effect for near-threshold stimuli (0 dB SPL, 2 kHz), that decayed rapidly with SPL. More specifically, the increase in the cumulative Ca_V_1.3 open probability (first detection method) was about 2.2 times higher using the stochastic receptor potential, and the average intra-cycle differences of the Ca_V_1.3 open probability (second detection method) were about 12 % higher. While these relative effects are significant, we acknowledge that the absolute differences in the open probability are rather minor and that near-threshold hearing may rely on other phenomena that go beyond the scope and resolution of our model.

### 2.9 Ribbon synapse

Excitation of the afferent type I auditory nerve fibers is governed by several consecutive steps – change of the IHC receptor potential, opening of Ca_V_1.3 channels in the IHC basolateral membrane, increase in the local Ca^2+^ concentration in the vicinity of the synapse, and ultimately a neurotransmitter release from fusing vesicles into the synaptic cleft.

The ribbon synapse holds synaptic vesicles close to the active zone, encoding the stimulus level in the rate of neurotransmitter release. The Ca_V_1.3 ion channels are expressed nearby the ribbon synapse, indicating a strong local effect of each channel. During the maturation period, channels are found progressively closer to the ribbon synapse (P. F. Vincent et al. 2018), suggesting that precise encoding requires co-localization of the ion channels and the ribbon synapse. It has been shown that the ribbon synapse itself can vary in size and shape (Moser et al. 2006), and it is thought that the morphology of the ribbon synapse plays an important role for its function (L.-Y. Wang and Augustine 2015). Specialization of the ribbon synapse and their long-term maturation has been confirmed recently by Niwa et al. (2021), who resolved differences in EPSC properties in pillar vs modiolar subgroups. Since the structure of the synapse can be experimentally resolved using transmission electron microscopy and super-resolution light microscopy (STED), see, e.g., Rutherford (2015), we designed the present model to reflect the spatial structure of the ribbon synapse.

The model is defined by positions of readily releasable vesicles and of individual Ca_V_1.3 channels. While the vesicle positions are experimentally directly accessible, resolving individual ion channels is currently not possible. However, the localization of ion channels with respect to the synaptic ribbon can already be measured (P. F. Vincent et al. 2018), and thus the function of the synapse may be studied with respect to variables like mean channel-vesicle distance. An example of a possible configuration of the ribbon synapse model can be seen in the Fig. 6C-E.

**Figure 6:**
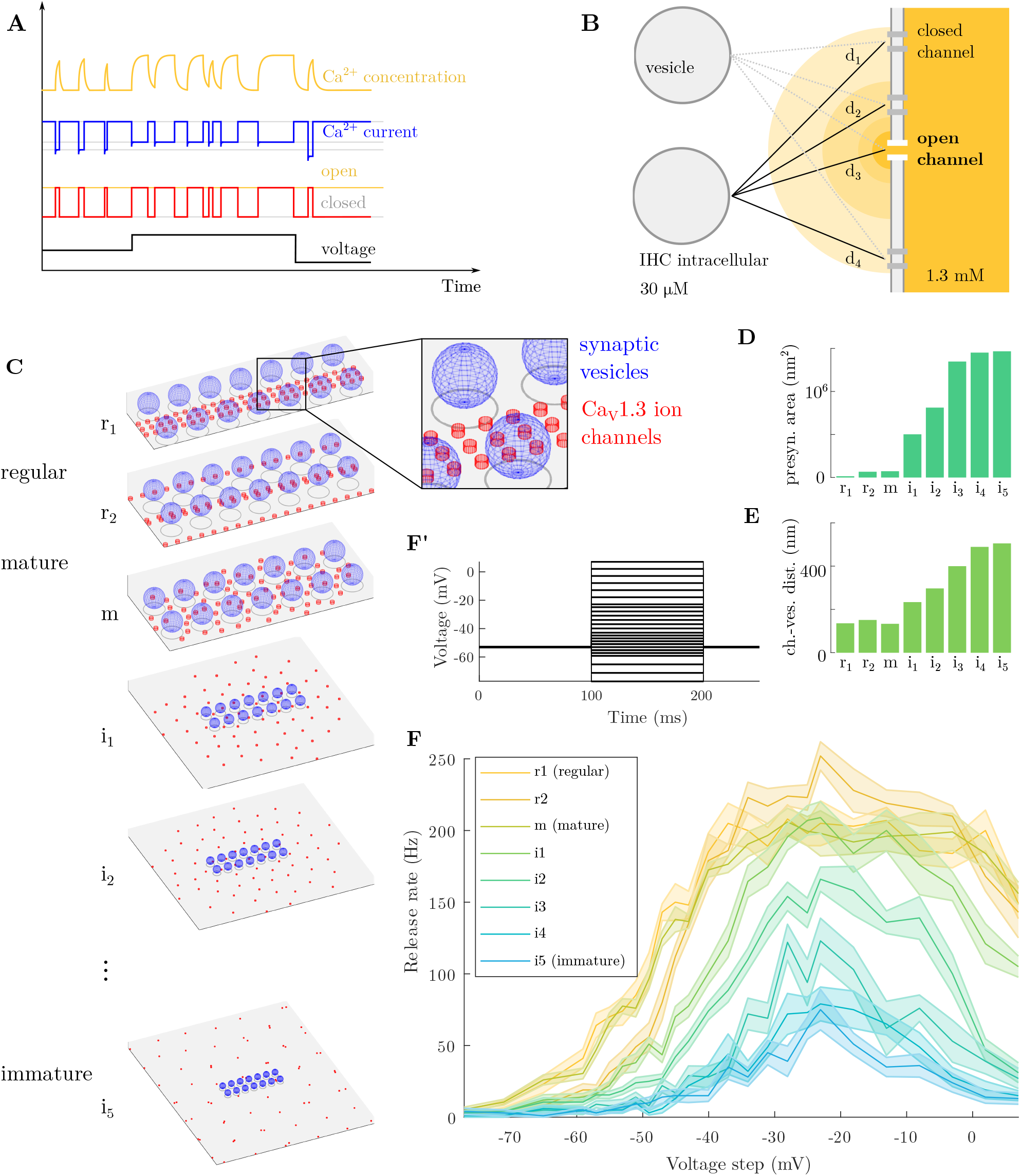
Effect of the morphology of the ribbon synapse. **A**: Illustration of a single Ca_V_1.3 stochastic opening. Depolarization increases the channel open probability, opening of the channel causes ionic current and concentration increase in its vicinity. **B**: Cumulative effect of many ion channels has to be taken into account, considering their positions. **C**: Sample models of the ribbon synapse, from a regular organization of the Ca_V_1.3 ion channels (r_1_, r_2_), through tightly-packed unorganized channels (m), up to spread-out configuration (i_1_-i_5_). **D, E**: Quantitative description of the channel distribution in terms of the presynaptic area (D) and the mean channel–vesicle distance (E). **F**: Average synaptic vesicle release rate for different channel configurations as a function of the peak voltage step in (F’).

The voltage sensitive Ca_V_1.3 channels play an important role in regulating the vesicle exocytosis and have been, therefore, the topic of intensive research. The crucial voltage-dependent steady state open probability has been reported repeatedly (Johnson and Marcotti 2008; Ohn et al. 2016; Zampini, Johnson, Franz, Knipper, et al. 2014; Zampini, Johnson, Franz, Lawrence, et al. 2010). While the results differ in absolute values of the curves (see Fig. M-3), they all agree that the open probability is very low close to the resting membrane potential of the IHC. Considering the total number of channels per active zone (80–600), it is clear that a release of a synaptic vesicle can be triggered by the opening of one or only a few channels (e.g. Magistretti et al. (2015)). Thus, compared to other models, each channel is explicitly calculated here.

Elementary properties of isolated Ca_V_1.3 channels have been investigated using patch-clamp techniques (Zampini, Johnson, Franz, Knipper, et al. 2014), yielding not only the voltage-dependent steady state open probabilities, but also the mean open/close durations. This allows us to express the open probability of the channel as a transfer rate within a two state Markov model.

Finally, one should take into account that that the channel activity can be temporarily blocked. Recent experimental findings indicate that this may be caused by protons released into the synaptic cleft alongside the glutamate neurotransmitter, particularly during multi-vesicular release events (P. F. Vincent et al. 2018).

Here, we describe the model of the ribbon synapse as a chain of events. After a channel opens, Ca^2+^ ions enter the cell following the concentration and voltage gradient. As a first approximation, the ionic current follows the Ohm’s law, the voltage being described by the Nernst equation. However, the current can be calculated more precisely by the Goldman–Hodgkin–Katz (GHK) equation, as is done in the present model. In addition, the immediate intracellular concentration in the channel vicinity is taken into account, with the extracellular concentration considered constant.

The Ca^2+^ concentration in the vicinity of a neurotransmitter vesicle is estimated considering all individual channels within the synaptic domain using a point-source diffusion model with space restricted by the membrane. The concentration at an arbitrary distance from the group of channels can be calculated analytically from the ionic current through each channel using the experimentally accessible diffusion coefficient of Ca^2+^ ions in cytoplasm. Compared to previous models, no ad hoc time-constants or any other calibration parameters are needed. In this way, the present model can predict differences in behavior of ribbon synapses of different spatial conformations.

The molecular mechanism behind Ca^2+^ governed synaptic vesicle exocytosis is still not fully understood, although considerable progress has been made in recent years, for example identifying Otoferlin as the Ca^2+^ sensor for both vesicle fusion and vesicle pool replenishment (Michalski et al. 2017). From a macroscopic point of view, the calcium-mediated exocytosis exhibits a highly positively cooperative influence of calcium, with the Hill’s coefficient for calcium dependent vesicle release from the Ribbon synapse being around five (Beutner et al. 2001; Jarsky et al. 2010), which is reflected in the present model.

Once a vesicle is released, its place is not immediately occupied by a new one, hence a vesicle pool depletion must be taken into account. Following Meddis (1986) and Sumner et al. (2002), we model the ribbon synapse as pools of free (i.e., ready to be released), re-processed, released, and freshly manufactured vesicles. In comparison to previous models, the release probability is calculated here individually for each free vesicle, depending on the respective local Ca^2+^ concentration. Additionally, each vesicle release is usually governed by only a small group of adjacent ion channels, compared to the previously published models, where individual channels have not been considered (instead, channels were modelled as a large population in a mean way). However, the number of “controlling” ion channels per vesicle is not a model parameter. Instead, the contribution of each channel is taken into account and, therefore, depending on the localization of calcium ion channels with respect to the ribbon synapse or depending on the stimulus level, the vesicle release can be caused both by a nearby channel or by a cumulative effect of multiple more distant channels. Finally, all vesicles from a single ribbon synapse released into the synaptic cleft are considered to be equal and the neurotransmitter re-uptake, vesicle factory, and reprocessing stores are joined into a single pool.

We have used the model to illustrate the effect of the post-natal maturation of the ribbon synapse (described by P. F. Vincent et al. (2018)) on its function. First, we have created different configurations of the ribbon synapse, having the same number of channels, but different presynaptic area and channel-vesicle distance distribution (Fig. 6C-E). Then, we simulated the response of the synapse to a voltage step in transmembrane potential and measured the mean vesicle release rate (Fig. 6F). Our results confirm that the “mature synapse”, i.e. the synapse with close colocalization of the vesicles and Ca_V_1.3 ion channels, has significantly increased sensitivity (especially close to the resting potential of the hair cell) compared to the “immature synapse” with high effective presynaptic area.

### 2.10 Auditory Nerve Model

An action potential is generated in the auditory nerve (AN) when a neurotransmitter is released from a vesicle at the ribbon synapse. However, when a delay between two such events is too short, only a single action potential occurs, due to refractoriness of the nerve fiber. Also, an action potential may not occur if the amount of neurotransmitter released is low and the resulting post-synaptic current does not reach the required postsynaptic activation threshold. Finally, spontaneous action potentials may occur due to channel fluctuations. To account for all these effects, we include an up-to-date implementation of the Hodgkin–Huxley model of the auditory nerve by Negm and Bruce (2014) as the last stage in the system. The model describes a single node of Ranvier in a mammalian auditory nerve fiber. The model contains four types of voltage-gated ion channels (fast sodium, delayed-rectifier potassium, low-threshold potassium and hyperpolarization-activated cation channels), and a passive leakage channel.

In order to account for stochastic phenomena in spike timing and threshold fluctuations, we used a stochastic version of the model including nondeterministic sodium channels. Specifically we use the Chow and White (1996) algorithm which describes the channel kinetics exactly with a finite-state Markov process. Other methods (deterministic Hodgkin-Huxley and approximate stochastic methods Fox and Lu (1994), which employs stochastic differential equations to approximate the Markov process) are also implemented (Bruce 2009) and available for specific use cases, where computational performance may be of concern.

To illustrate the functionality of the final stage of the present model, we performed simulations of the auditory nerve spiking for a single synapse morphology using 4 kHz pure tones of varying sound levels, reconstructing thus the input-output curve of a nerve fiber at its characteristic frequency. The results are presented in the Figure 7. We analyzed the basic spiking characteristics in terms of a post-stimulus time histograms (PSTH) and highlight here the observed adaptation, i.e., different probability of activation at the onset of the stimulus compared to the steady state situation. The present firing patterns (Fig. 7A) can be classified as a medium spontaneous rate (SR) fibre, with mean-rate detection threshold at about 30 dB SPL, that saturates (in a steady state) at approximately three-times its SR. It shows the typical experimentally observed onset-peak and post-offset-decrease of firing rate, both being more prominent with SPL. The steady-state rate-level function (Fig. 7C) saturates for high sound level (in this case 70–80 dB SPL), the onset rate-level function does not saturate in the explored range, but shows a decreased growth speed above 80 dB SPL, which is in agreement with experimental evidence (Smith and Brachman 1980).

**Figure 7:**
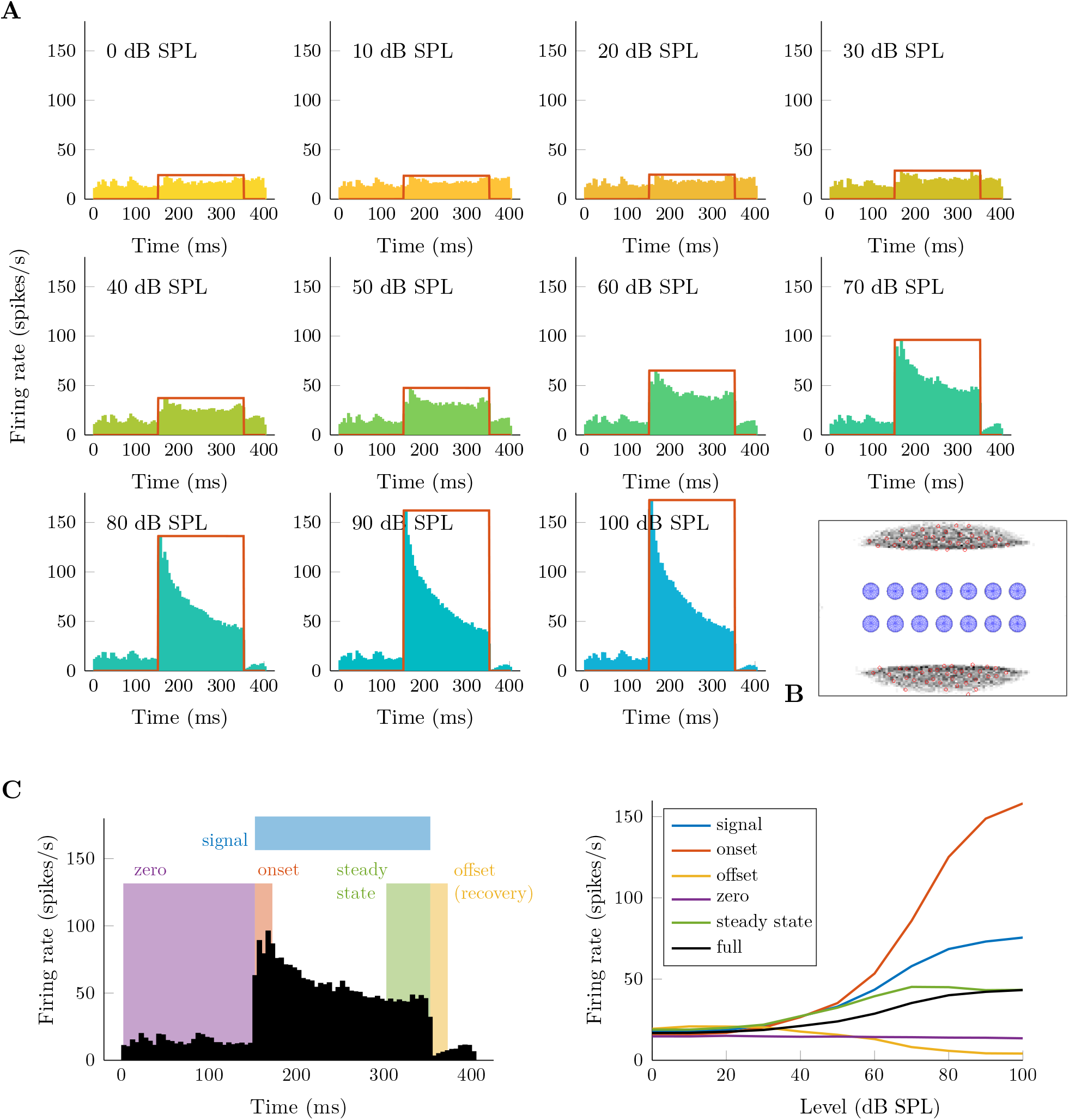
Auditory nerve response: **A**: Response of the auditory nerve (AN) to 4 kHz tones of 0 to 100 dB SPL in the form of post-stimulus time histograms (PSTH). The depicted firing rate was averaged over 54 independent simulations of a single ribbon synapse and 100 independent simulations of AN firing (per each synapse simulation) with a 5 ms bin width. The 200 ms stimulus was preceded and followed by 150 and 50 ms of silence; its amplitude is shown by a red line. **B**: Morphology of the synapse, view perpendicular to the cell membrane; dark areas reflect regions with high probability of Ca_V_1.3 expression. **C**: Rate-level functions (right) derived from simulated AN nerve responses, decomposed according to the scheme (left) into onset, steady state (during stimulus), offset (recovery), and zero (steady state without stimulus). As expected, the “onset” shows the steepest slope, while the steady state saturates above approx. 60 dB SPL. Interestingly, the “notch” that can be observed in the PSTHs of high-level tones after the stimulus fades (panel A) is more pronounced with the sound level, which can be observed by a decrease of the “offset”-domain rate-level function with SPL. The notch, however, originates in lower levels of the present model as can be seen in the Figure S-4 and is caused by the potassium channel kinetics.

The employed synapse morphology is presented in the Figure 7B. Any synapse configuration can be simulated and the effect of the morphology is analyzed in terms of the AN spiking characteristics. Here, we analyzed the effects of the Ca_V_1.3 channel distribution using the example of synapse maturation, with further analysis that goes beyond the scope of this paper being feasible.

### 2.11 The integrated model

The present approach to the computer model of the peripheral auditory system allowed us to extend the focus from a single cross-section to an array of 3000 coupled cross-sections (each containing a single IHC and four joined OHCs), explicitly calculating the response of each of them. Furthermore, for each IHC tens of individual ribbon synapses can be modelled individually. Being formulated and numerically solved in the time domain, the model can simulate the response to an arbitrary acoustic signal. As an illustration of the different model outputs, we show here the responses to a single word “greasy”, and a noise-band bursts, see Fig. 8.

**Figure 8:**
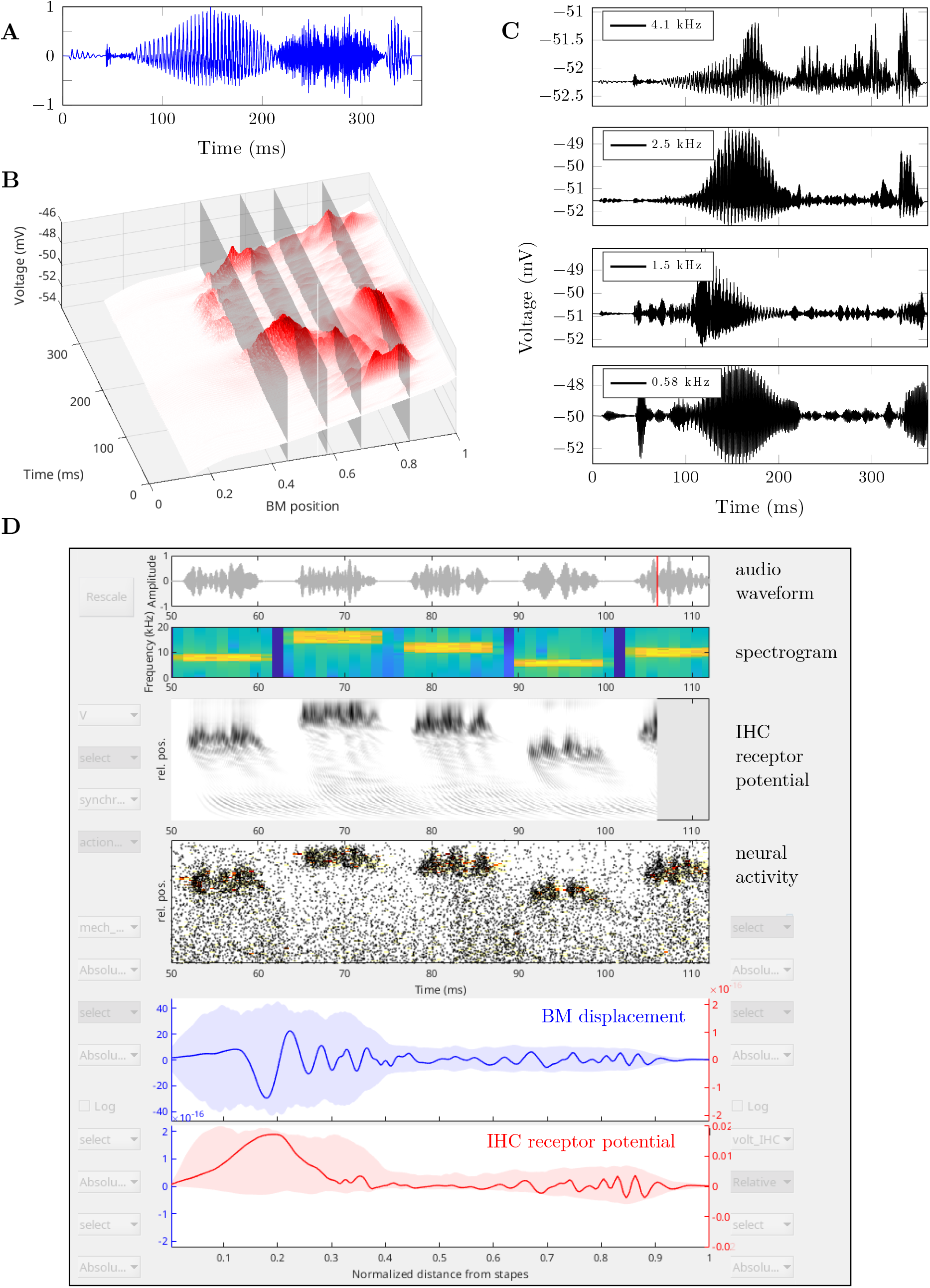
Output analysis and the graphical interface: **A-C**: Example model output of a simulation of person saying the word “greasy”. **A**: Stimulus waveform. **B**: IHC basolateral voltage as a function of time along the tonotopy axis. **C**: Cuts of the three-dimensional graphs in B showing the response at specific characteristic frequency locations. **D**: Snapshot from the graphical user interface (GUI) allowing the user to visualize any calculated variable on the fly as well as after the simulation has been completed. Videos of different simulations are available as part of the supplementary material available online at doi.org/10.6084/m9.figshare.20348271.

Thanks to an efficient implementation, we are able to simulate hundreds of different ∼100 ms signals in an over-night run on a high-end personal computer, so in general, the work with the present model can be realized in an interactive way without the need for high performance computing systems. Importantly, the modular implementation allows for easy extensions of the computer program, both in terms of new features and specific analyses.

## 3 Discussion

The present model is designed with a focus on detail and a bottom-up approach, based on the appropriate physical laws and principles. Its principal advantage lies in its ability to examine the impact of all the factors included in its design, ranging from the mechanical-to-electrical transduction, over the electrical properties of ion channels in hair cells, to the morphological characteristics of the synapse. Our model was built upon a foundation of older models (Bruce 2009; Lopez-Poveda and Eustaquio-Martín 2006; Mammano and Nobili 1993; Meddis 1986; Mistrík, Mullaley, et al. 2008; Negm and Bruce 2014; Pascal et al. 1998; Strelioff 1973; Sumner et al. 2002; Vetešník and Gummer 2012), with other ones being developed in parallel (e.g. Heil et al. (2011), Motallebzadeh et al. (2018), Ramamoorthy et al. (2007), Sasmal and Grosh (2019), and Verhulst, Altoè, et al. (2018)). Here, we briefly discuss these models putting the present one in the context of the existing literature, focusing on the cochlear mechanics and ion flows therein and, in particular, on the interface between the hair cells and the axons of the auditory nerve, including the ribbon synapse.

In the field of modelling of the cochlear mechanics several approaches have been employed in parallel, which differ by the level of detail of description (and the corresponding computational costs), ranging from digital filter-bank models to finite element 3-dimensional models (Motallebzadeh et al. 2018; Sasmal and Grosh 2019). In the present study, we have built upon an older model of intermediate complexity originated by Mammano and Nobili (1993) and Nobili and Mammano (1996) who showed that such a model, where the cochlear amplifier is explicitly based in the OHCs, can reproduce a wide range of phenomena, such as the two-tone suppression. The model is 2-dimensional, taking into account the height of the cochlear duct, which has been recently been shown to be important for a proper BM response (Altoè and Shera 2020). Vetešník and Gummer (2012) improved the original model, calibrated it to the human cochlea, and along with Vencovský et al. (2019) employed it to the study of distortion products and otoacoustic emissions.

As far as ionic flows in the cochlea are concerned, the original model by Strelioff (1973) included all three scalae, but the organ of Corti was oversimplified to a single resistor and a single voltage source per cross-section. Later, Mistrík, Mullaley, et al. (2008) adopted the general idea of a three-dimensional electric model of the entire cochlea, allowing to investigate the ionic flow not only within a single cross-section, but also in-between them along the tonotopy axis. Focusing on the organ of Corti they were also able to model the ionic currents through both inner and outer hair cells. However, the basolateral hair cell conductance was considered to be constant (i.e., voltage independent), the channel kinetics were not accounted for, and the channel subtypes with different activation properties were not resolved, in contrast to what has been done for a single IHC by Lopez-Poveda and Eustaquio-Martín (2006) and later by Dierich et al. (2020). The tonotopic gradient in the properties of the hair cells was not included in a more recent model by Mistrík and Ashmore (2010) either. Moreover, the nonlinear capacitance, specific to outer hair cells, has not been included in any of these models. The present approach overcomes these limitations and models each of the 3000 cochlear partitions with a separate set of parameters.

For the IHC synapse, it was Meddis (1986) who laid grounds to the modeling approach and his concept of synaptic neurotransmitter “stores” has been in varying forms adopted also in more recent studies including the present one. Later, Sumner et al. (2002) introduced stochasticity of the synaptic vesicle release, which served as inspiration for the present work. While Sumner et al. (2002) were pioneers at aiming to provide a model of the whole auditory periphery, their model components were rather simplified (this concerns, in particular, the filter-based model of cochlear mechanics). Additionally, the calcium currents and concentrations in the vicinity of the ribbon synapse were not modelled in sufficient detail, and the model was calibrated only to a single tonotopic location of 18 kHz. In contrast, the present model is tonotopically resolved and detailed enough to be able to capture even subtle effects, such as the post-natal maturation of the ribbon synapse.

## 4 Conclusions

In conclusion, we have built a complete computational model of the human peripheral auditory pathway from the outer ear to the afferent auditory nerve. While most of the cochlear models previously published target only a specific part of the auditory pathway, here we present a complete model of the peripheral auditory system built in a bottom-up way in a physically and physiologically well justified way and with parameters reflecting up-to-date experiments. Within such a cascade approach, where the output from each stage of the model serves as input to the next stage, it has been our goal that the output from each model stage accounts for as many aspects of corresponding physiological responses as possible. On the one side, the present model represents a research tool that allows to analyze the effects of the chosen parameter values and investigate the consequences of changes thereof. In this way, it provides a deep quantitative insight into the mechanism of hearing. On the other side, various forms of hearing impairment can be readily implemented and their effects investigated. As one such example for the cochlear model, effects connexin mutations on hearing impairment can be modelled in significantly more detail than attempted previously (Mistrík and Ashmore 2010).

The model aims also to serve as an in-silico diagnostic tool to identify pathological mechanisms underlying different forms of deafness. In particular, the present model can predict the effect of disorders originating in any part along the peripheral auditory pathway, in particular in the mechano-electric transduction, in the electro-mechanic response of the OHCs, in the neurotransmitter release in the IHCs, as well as in the action potential generation in the primary auditory nerve. In this way, the model can help to diagnose a hidden hearing loss and possibly also distinguish between its main causes, which is imperative in predicting the viability of implantation with a cochlear implant (Shearer and Hansen 2019). Due to its sufficiently detailed level of description, the model is also able to determine the effect of new mutations in genes responsible for hearing loss (Dror and Avraham 2010). In addition, in conjunction with experimental measurements, the model can serve as a tool to evaluate the effect of potential drugs targeting deafness (Isherwood et al. 2022). Finally, a straightforward application of the model lies in the investigation of the functionality of cochlear implants. The model can be straightforwardly extended to capture for example the combined effect of using the implant together with hearing aids, which will be instrumental in developing new simulation and fitting strategies for cochlear implants.

## 5 Methods

### 5.1 Outer ear filtering

The outer ear transfer characteristics are implemented via a set of independent band-pass filters as proposed by Meddis (2011). Specifically, we used *N* = 11 fourth-order Butterworth band-pass filters, that transform the sound pressure *p*_0_ to the pressure at the tympanic membrane *p*_tymp_:

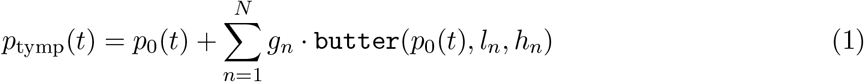

where *g*_*n*_, *l*_*n*_, and *h*_*n*_ are parameters of the filter – gain, low- and high-cutoff frequencies. Note that even though this design is purely additive, moderate attenuation at certain frequency bands can be achieved because of phase shift introduced by the filters, see Figure 2A.

### 5.2 Middle ear lumped-element model

The middle ear is represented as a lumped-element model of selected structures — namely the eardrum, the middle ear cavities, malleus, incus, stapes, the cochlea — and by their connections. The design has been taken directly from Pascal et al. (1998). For simplicity, we chose to omit the non-linearities in the response caused by the acoustic reflex and the influence of the angular ligament on the maximum stapes displacement, which appear for very high-level stimuli.

The mechanical model can be translated into an equivalent electrical representationand be driven by a voltage *V*_0_ and the response can be observed in the form of voltage across the ‘cochlea’ resistor, *V*_cochlea_. The values of *V*_0_ and *V*_cochlea_ can be related to the tympanic membrane pressure *p*_tymp_ and the oval window pressure *p*_cochlea_ as

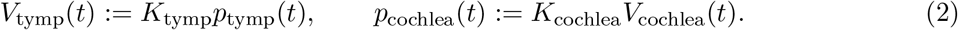

Since the model is linear in respect to the driving voltage (and therefore the acoustic pressure), the conversion constant *K*_tymp_ can be set to 1 V/Pa. The scaling constant *K*_cochlea_ [Pa/V] needs to be calibrated for the model to yield correct oval window pressure levels.

The electrical circuit representing the middle ear can be translated to a mathematical description using the Modified Nodal Analysis (MNA). This method results in a differential-algebraic equation (DAE) of the form

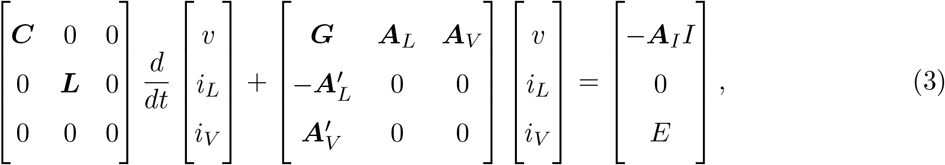

where ***C, L*** and ***G*** are matrices defined by the capacitances, inductances and conductances of the circuit elements, ***A***_*L*_, ***A***_*V*_, and ***A***_*I*_ are defined by their connectivities, *I* = 0 and *E* = (*V*_tymp_, 0, …, 0) are current and voltage sources and *v, i*_*L*_ and *i*_*V*_ are the node voltages and currents through inductors and voltage sources. The equation can be solved in the time domain as well as the frequency domain by employing the Fourier transform. Here, we used routines based on the symbolic circuit solver by Cheever (2018) to assemble the matrices and solve the system in the time domain using standard DAE integrators (MATLAB 2021).

In their original paper Pascal et al. (1998) shown that such a model can be calibrated to reproduce an experimental sound pressure-intracochlear fluid pressure transfer function of a single individual. Since the cochlear mechanics model we employ in the later stage requires the stapes acceleration as an input, we recalibrated the model to a recent experimental tympanic membrane pressure-stapes velocity average data by Chien et al. (2009), see Figure 2B.

### 5.3 Cochlear Mechanics

Cochlear mechanics is governed by two partial differential equations for functions *ξ*(*t, x*) and *η*(*t, x*) that represent the displacement of the basilar membrane and the deflection of OHC stereocilia. The variable *x* here and in all future occurrences describes the position, along the longitudinal axis of an “uncoiled” cochlea, normalized so that *x* = 0 at the base and *x* = 1 at the apex; the variable *t* means time in all occasions.

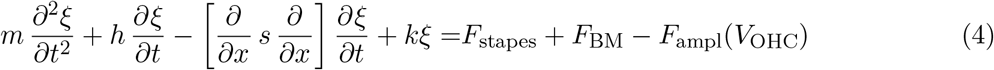

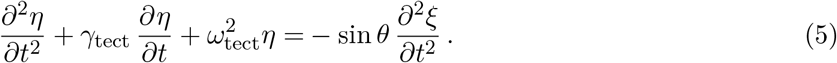

The equations are parameterized by the mechanical properties of the BM such as the mass *m*(*x*), damping *h*(*x*), shearing resistance *∂*_*x*_*s*(*x*)*∂*_*x*_, and stiffness *k*(*x*), and tuning characteristics of the 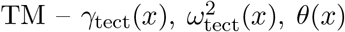. The model is driven by the stapes motion *σ*(*t*) (computed by the model of middle ear) via a fluid coupling between the stapes displacement and the BM displacement

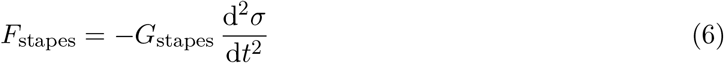

modeled by the Green’s function *G*_stapes_(*x*). Individual BM segments are also connected via fluid coupling

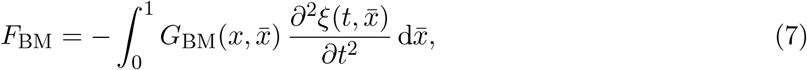

where 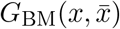 is the Green’s function. Here we refer the reader to the original paper by Vetešník and Gummer (2012) for detailed description of the parameters.

The differential equations are coupled to the model of ionic currents through the cochlea by regulating the conductance of MET channels (described later) and via the OHC transmembrane voltage *V*_OHC_, that induces an active force on the system resulting in amplification. The amplifier force *F*_ampl_ is explicitly computed (for each position *x*) via the differential equation

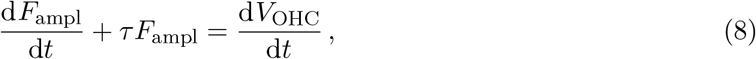

which is a high-pass filter defined by the time constant *τ*. The force *F*_ampl_ needs to be zero at steady state, which is fulfilled for this equation by choosing a proper initial conditions. The filtering inherently involves a magnitude change and phase shift, however, for small enough *τ*, the filtering affects only very low frequency components, leaving the physiologically relevant unchanged (see Supplementary Fig. S-7).

Due to the novel description employing directly the OHC receptor potential, there is a (nonlinear) feedback loop between the model of cochlear mechanics and the ionic currents. Consequently, the partial differential equation 4 has to be solved simultaneously with the differential-algebraic equation 12.

### 5.4 Channel dynamics

Measuring the basolateral K^+^ currents in isolated guinea-pig IHCs, Kros and Crawford (1990) determined that each of the channel subtypes exhibits kinetics with two closed states (*C*_1_, *C*_2_) and one open state (*O*) governed by voltage dependent transfer rates *α, β, γ*, and *η*

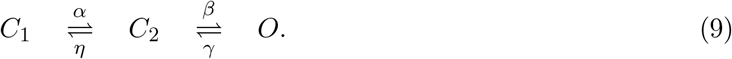

Considering a population of the channels, the stochastic dynamics of the channels can be described in a mean way by differential equations, see Lopez-Poveda and Eustaquio-Martín (2006) for derivation and details. They show that open state probability *O*(*t, V*) of the channels can be equivalently described by a second-order differential equation

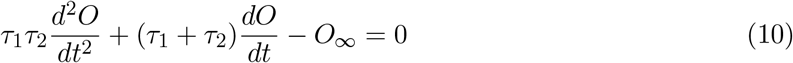

where *τ*_1_(*V*) and *τ*_2_(*V*) are voltage dependent time constants and *O*_*∞*_(*V*) is voltage dependent steady-state open probability. The time constants *τ*_1_, *τ*_2_ and the steady state open probability *O*_*∞*_ are directly related to the transfer rates from Eq. 9, while being experimentally accessible. We have applied the methodology for both IHCs and OHCs; an independent set of these nonlinear differential equations, one per channel subtype and per hair cell, is inherently coupled with the system 12 describing the ionic currents through the cochlea, as the instantaneous channel conductance is proportional to the open probability:

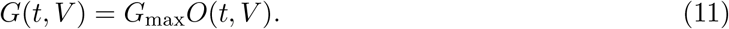

This creates yet another nonlinear feedback loop in the system, and therefore the Eq. 10 has to be solved concurrently with the Eq. 12.

### 5.5 Model of a single hair cell

Electrophysiological experiments are usually performed on isolated hair cells. The hair bundles are then bathed in the same extracellular solution as the rest of the cell, eliminating the endocochlear potential. In order to compare directly to in-vitro experiments, we implemented a model of an isolated hair cell that can simulate whole-cell patch clamp recordings as well as response to hair bundle stimulation.

The model (Fig. 3E) is designed in a similar fashion as the model of a cochlear cross-section. It does not include the endocochlear potential, but is driven as in a standard patch-clamp experiment by a current or voltage pulses. Due to it’s computational simplicity, it can be used for parameter estimation using multivariate optimizers (we have used differential evolution). Results of optimization can be seen in the Figure 3F-J.

### 5.6 Equivalent circuit of the cochlea

In addition to resistors and capacitors representing the conductive properties of cellular membranes expressing ion channels or prestin, the circuit is composed also of voltage sources accounting for actively maintained electric fields and ionic concentration gradients across the cellular membrane through the Nernst equation. Since the primary interest is in the current flow through the sensory cells of the cochlea, the inner and outer hair cells are implemented in the most detail (apical and basolateral membranes). An equivalent electrical circuit of a single cross-section of the cochlear partition can be seen in the Figure 3A. The outer hair cells are represented at each cross-section by a single element with its values scaled up accordingly. Since it is assumed that the outer hair cells within one cross-section are independent and identical, this does not change the results.

The construction of the equivalent electrical circuit representing the whole cochlear partition is concluded by a longitudinal connection of the partial “cross-section” circuits. As seen in the Figure 3A, the cross-section circuits are linked together by groups of resistors, representing corresponding sections of conductive paths in the scalae (SV, SM, ST), and in the stria vascularis (StV), spiral limbus (SL) and organ of Corti (OC). In the full circuit, there is one cross-section sub-circuit for every row of hair cells (single IHC and four OHCs), resulting in approximately 3000 sections for the human cochlea. For some experiments, due to computational requirements, some of these sections were lumped together while adjusting the values of the electric elements. This can be done without significant loss in accuracy; one only has to consider the loss of spatial resolution along the longitudinal axis of the cochlea.

Since the circuit only contains passive linear elements (resistors and capacitors) plus only independent voltage sources are considered, the modified nodal analysis (MNA) can be used for constructing the circuit equations. The resulting system of differential equations can be written as

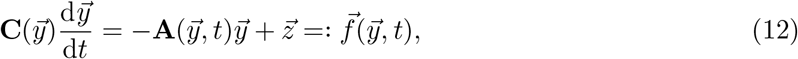

where the unknown vector 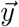 holds the voltage at nodes of the circuit and current through voltage sources, the matrix 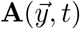 is a time-, voltage-, and current-dependent block matrix defined by the interconnections of resistors and voltage sources, 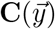 is a singular matrix defined by the capacitors, the vector 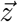 contains voltage and current sources. Since the matrix 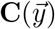 is singular, the system 12 is in fact a differential-algebraic equation (DAE), and consequently a suitable numerical method must be employed.

Let us now describe the equation and its connection to the cochlear model in more detail. In the following, let *n* denote the number of nodes (except for the ground node *zero*) in the circuit and *m* the number of independent voltage sources.

The matrix 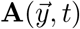 is generally constructed as a block matrix

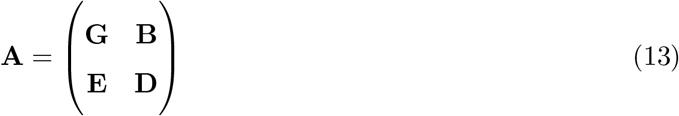

where **G** ∈ ℝ^*n,n*^ holds the conductances of the circuit elements between the nodes, matrices **B** ∈ ℝ^*n,m*^ and **E** ∈ ℝ^*m,n*^ are determined by connection of the voltage sources, and **D** ∈ ℝ^*m,m*^ is zero if only independent voltage sources are considered.

The vector 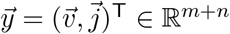 contains unknown quantities — voltages at nodes 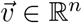 and currents through voltage sources 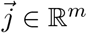. The vector 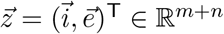 is defined by current and voltage sources values — the sum of all current sources at each node defines the vector 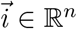, the electromotive force of each voltage source defines the vector 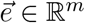.

An equivalent electrical circuit consisting of *k* cross-section each containing *n* nodes and *m* voltage sources results in a system of *k ×* (*m* + *n*) unknowns. In the current design, (*m* + *n*) = 17, and *k* can vary from 600 up to 3000. Consequently, the computational requirements are rather high. Since a significant portion of the resources was spent on creating the time-dependent matrix **A**(*t*) during the construction of the Jacobian matrix or while computing the right-hand side of the (12), an alternative to the classic MNA procedure was employed. It exploits the fact that only the submatrix **G** of the matrix **A** is time and voltage dependent and, moreover, that **G** can be expressed as

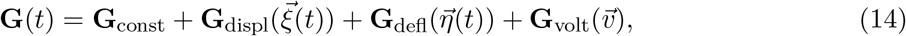

where 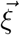 is the BM displacement, 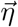 is the deflection of the stereocilia in OHCs. Moreover, the voltage-dependent part of **G** can be further decomposed to

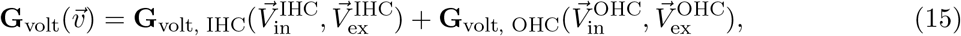

where 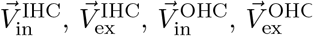 are IHC and OHC intra- and extracellular voltages.

The Jacobian matrix of the DAE is defined as

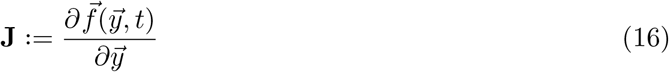

where 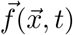 is defined by the system

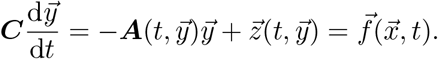

The Jacobian matrix is necessary for most DAE solvers, including the employed matlab ode15s.

If 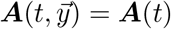, then 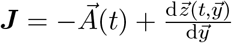, otherwise product rule must be used

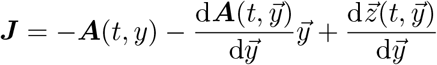

Note that 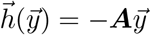 is matrix times vector, i.e.

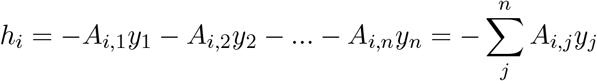

and therefore

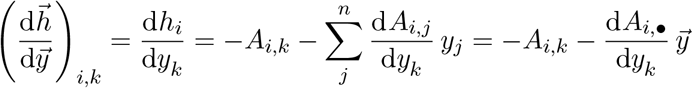

where *A*_*i,•*_ is a *i*-th row vector.

**Figure M-1:**
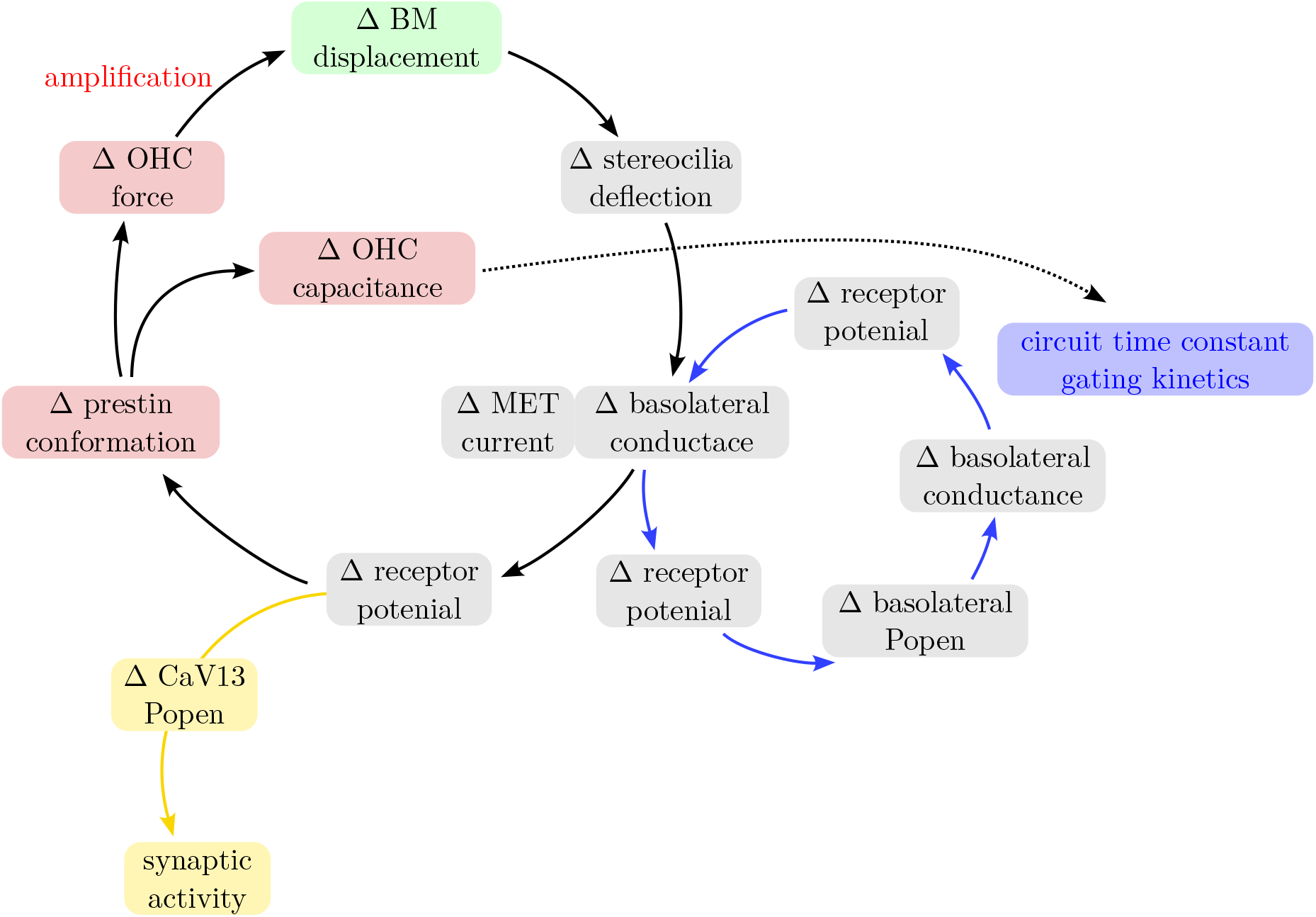
Feedback loops in the model of cochlear mechanics and ionic currents.

### 5.7 Closed feedback loop in the system of models

Deflection of the OHC stereocilia *η* changes the instantaneous conductance of the MET channels 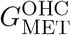 changing the potential across the apical, and also the basolateral OHC membrane *V*. The voltage gated basolateral channels adjust their open-state probability *O* and conductance 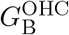, pushing the voltage change partially back towards the equilibrium. At the same time, the voltage gated prestin protein changes its conformation and inflicts a force on the basilar membrane. If this happens in phase with the BM oscillation, it results in amplification of the motion of the coupled BM–TM system. Finally, the simultaneous motion of the BM and TM induces a deflection of the OHC stereocilia, thus closing the loop. The feedback is visualized schematically in the Figure M-1 and also through the Jacobian matrix of the system in the figure M-2.

### 5.8 Ribbon Synapse

The ribbon synapse is defined by the positions of readily releasable (membrane-tethered) vesicles and the Ca_V_1.3 ion channels. Vesicles attached to the ribbon further away from the membrane and cytosolic vesicles are currently not considered. In the present configuration of the model, there are 14 readily releasable vesicles distributed in two uniform rows (Khimich et al. (2005) states 16–30 ready-to-release docked vesicles), covering an area of approx. 120 × 400 nm (approx. in line with Moser et al. (2006)). The vesicles in the model are considered identical spheres with a radius of 20 nm based on our measurements of electron micrograph (circular cross-sections with radius *r* = 16 *±* 2 nm from Frank et al. (2010), elliptical cross-sections with *a* = 25 *±* 4 nm, *b* = 17 *±* 2 nm from Rutherford (2015)). The vesicles are located 5 nm from the plasma membrane, again estimated from the micrographs. Note that the number of vesicles, their size and spatial distribution are intended as model parameters.

**Figure M-2:**
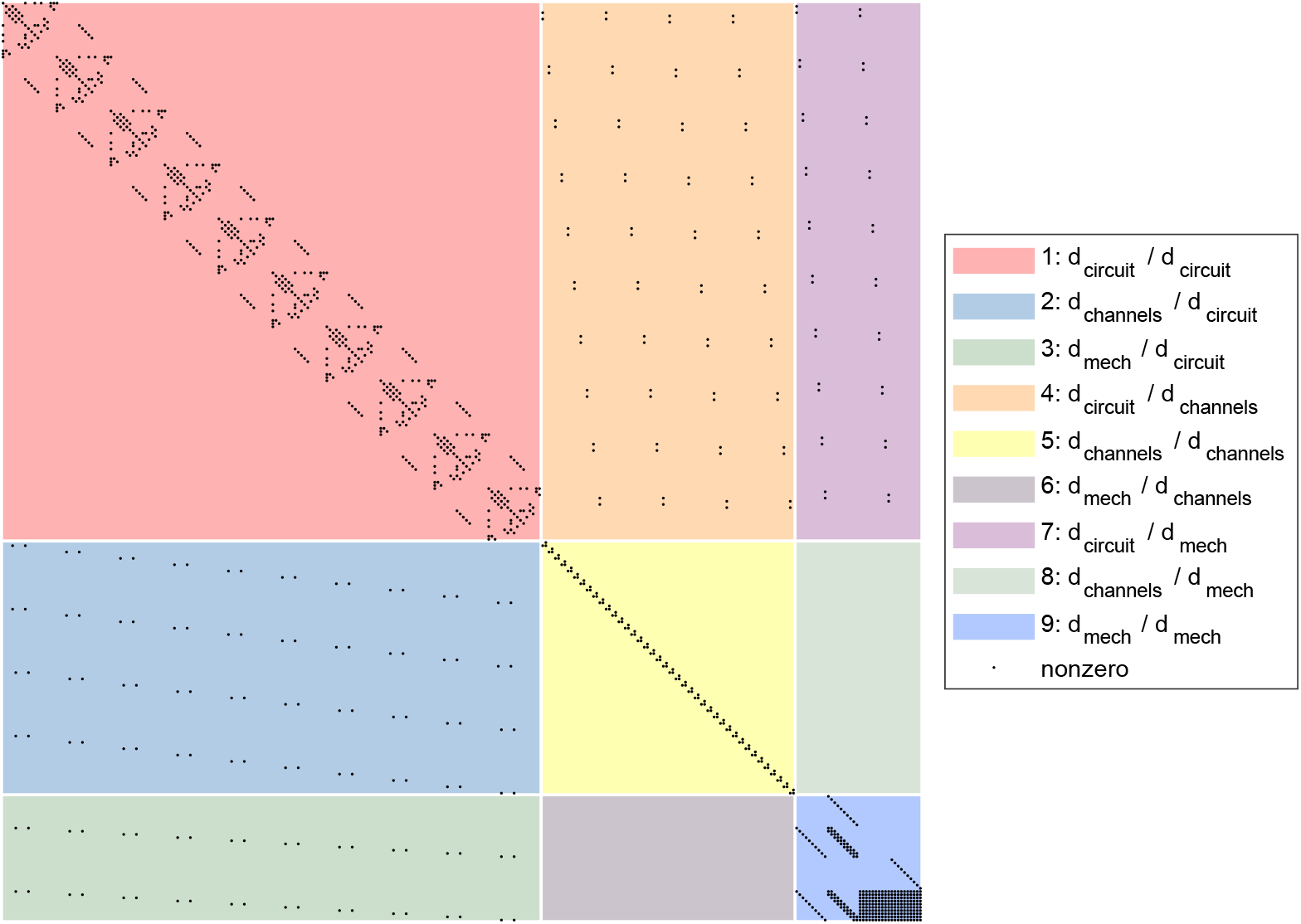
The interaction between the model blocks visualized via the Jacobian matrix. Internally, the model is composed of three blocks: the electrical circuit, the gating model of the basolateral IHC and OHC ion channels, and the mechanical model, each composed of multiple equations. When the *i*-th equation depends on the *j*-th component of the solution, the derivative *∂f*_*i*_/*∂y*_*j*_ is generally nonzero, and therefore, the *J*_*i,j*_ element of the jacobian matrix is nonzero. In the figure, nonzero elements are displayed as black dots. The matrix is shaded with colors corresponding to different classes of the dependence. The diagonal blocks represent internal dependence within the circuit, the channels and the mechanical models. The off-diagonal blocks instead symbolize the feedback between the models. The off-diagonal blue block (2) represents the voltage dependence of the basolateral IHC/OHC channels, the green block (3) shows the voltage-dependent OHC amplifier, the orange block (4) represents the basolateral IHC/OHC conductance change upon the opening of the channels, and finally the purple block (7) represents conductance change of the MET channels induced by vibrations of the BM and deflection of the stereocilia. Note that for visualization purposes, the number of cross-sections (corresponding to the size of the system) was reduced to 10. Otherwise, the fine structure of the nonzero elements would be hidden due to the image resolution.

The Ca_V_1.3 ion channels are represented by their positions on the membrane surface. The distribution of the ion channels in the active zone of the synapse is an input parameter of the model. For an ease of use, we have implemented a simple Monte Carlo simulation controlling the spread of the ion channels, producing random configurations from wide-spread-out to closely-packed (simulating synapse maturation) by changing just two parameters.

The number of ion channels in a IHC can be estimated experimentally via non-stationary fluctuation analysis of Ca^2+^ tail currents. For the mammalian IHC Brandt (2005) gives about 1700 Ca^2+^ channels, similar numbers have also been reported later (P. F. Y. Vincent et al. 2014; Wong et al. 2013). The channels are mostly expressed around the active zones (about 5–30 per IHC, Moser et al. (2006)). However, Magistretti et al. (2015) argues that the total number is in fact higher, when accounting for inactivated channels. They give 6400 channels (middle turn) and 1152 in about 16 active zones (basal turn). Along these lines, Zampini, Johnson, Franz, Lawrence, et al. (2010) gives 9600 channels (P5-P10 mouse, apex). Zampini, Johnson, Franz, Knipper, et al. (2014) gives about 16000 channels in about 21 active zones (adult gerbil, middle turn 2 kHz). A single channel conductance is estimated by Zampini, Johnson, Franz, Knipper, et al. (2014) to 15 pS.

The Ca_V_1.3 channels are known to be voltage-sensitive, the open probability is usually expressed in terms of a Boltzmann function

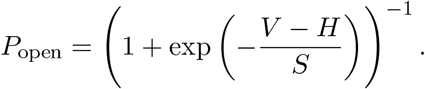

The open probability can be accessed indirectly by measuring the whole cell Ca2+ conductance or by analyzing open and close distributions of single channel measurements. Several reported Boltzmann curves can be found in Fig. M-3 as well as the curve used in the present model. It has been suggested that the parametrization of the Ca_V_1.3 open probability can change among individual active zones, displaying a pillar–modiolar gradient (Ohn et al. 2016), yielding a possible extension to this model.

Channels in the synapse model can be either closed or open, corresponding to two Markov states

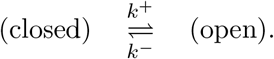

The transfer rates *k*^+^ and *k*^*−*^ can be expressed as

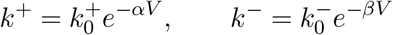

where *α* (unit V^*−*1^) and 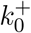 (unit s^*−*1^) are control parameters and

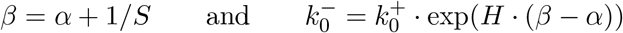

Then the probability transfer matrix is

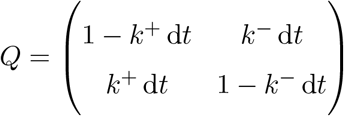

where d*t* is the simulation time step.

The control parameters *α* and 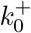 then control the timescale in which the transfers occur (the kinetics of the process). Their values can be obtained by comparing the mean open and closed times of the Markov process with experimental data. We have calibrated the rates to reflect the mean open and closed times of single channel patch-clamp measurements, as reported by Zampini, Johnson, Franz, Knipper, et al. (2014).

At any point in the space in the vicinity of the active zone, the Ca^2+^ concentration *C* is calculated as a sum of the background concentration *C*_bg_ and the partial concentrations *C*_*i*_ caused by elementary current through the *i*-th ion channel. The background (bulk) endolymph calcium concentration ranges from 17 µM (base) to 41 µM (apex) (Salt et al. 1989). For each location of interest and each ion channel, the concentration *C*_*i*_ is calculated via the single-point source diffusion equation

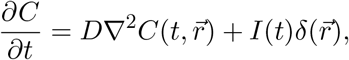

**Figure M-3:**
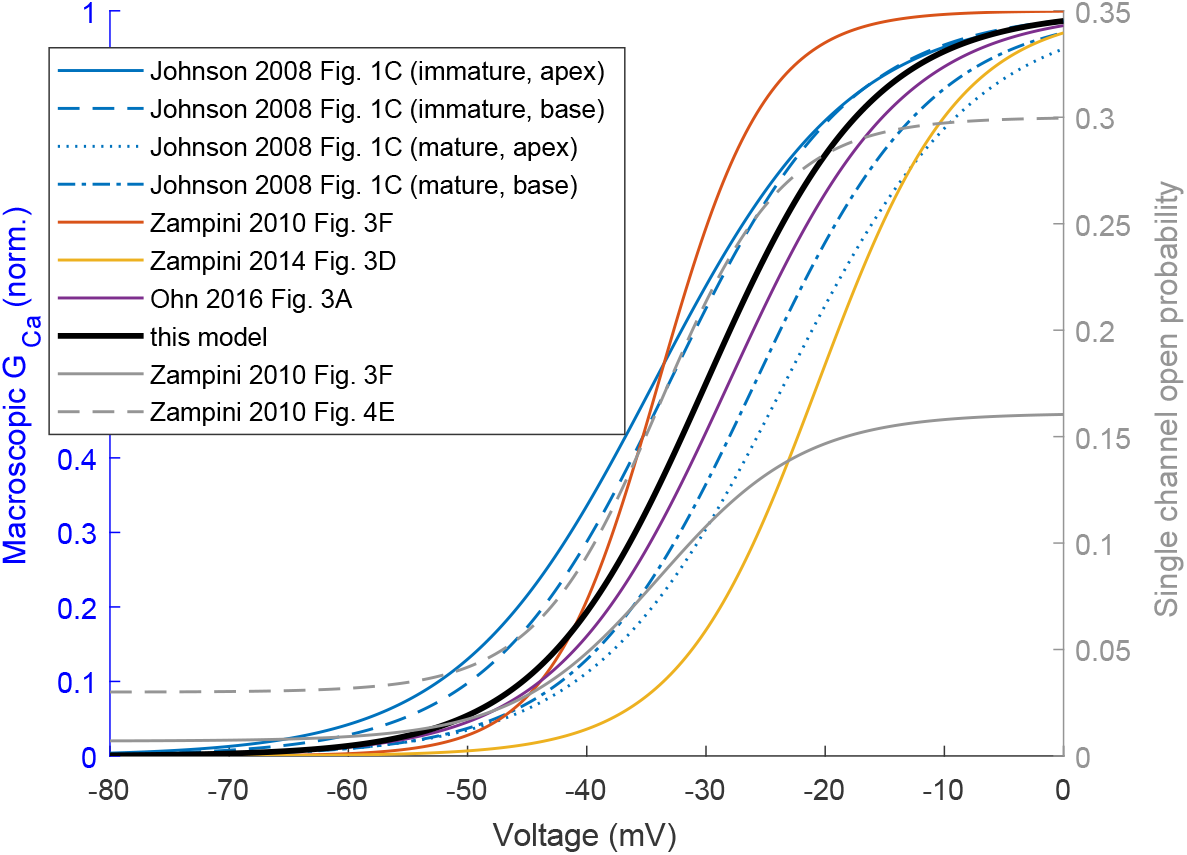
Ca_V_1.3 normalized conductance and open probabilities used in this model compared to available experimental data by Johnson and Marcotti (2008), Ohn et al. (2016), Zampini, Johnson, Franz, Knipper, et al. (2014), and Zampini, Johnson, Franz, Lawrence, et al. (2010).

where *r* is the distance from the ion channel, *D* is the diffusion coefficient, *I* current through the channel and *δ* is the Dirac delta function. We solve this equation analytically (using propagation integrals) for each time step, as described in Gillespie (2020). The main advantage of this method is that since the solution is exact, it does not require any spatial discretization and the time steps can be arbitrarily large, as long as the current through the channel can be considered constant.

For each vesicle, we are taking into account all Ca_V_1.3 channels within the active zone (and the distance from each channel.

The calcium current through individual channels can be calculated via the Goldman–Hodgkin–Katz equation

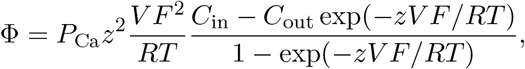

where *V* is the receptor potential, *P*_Ca_ is permeability, *R* is the gas constant, *F* is the Faraday constant, *z* = 2 is the calcium cation valence, *C*_in_ is the intracellular and *C*_out_ = 1.3 mM the extracellular calcium concentration (assumed constant along the tonotopy axis) (Salt et al. 1989).

It is unclear in which distance from the channel should one calculate the effective calcium reversal potential for that channel. Zampini, Johnson, Franz, Knipper, et al. (2014) measured macroscopic Ca^2+^ currents in middle-coil IHC bathed in high Na^+^, 5 mM Ca^2+^ solution in a voltage range *−*70 to 0 mV and estimated the calcium reversal potential to 28 *±* 2 mV. Having performed analogous simulations, we estimated the distance of 20 nm to yield similar reversal potentials for sustained Ca^2+^ currents.

The probability of vesicle release is proportional to the calcium concentration at the calcium sensor, saturating for high concentration values, which is achieved for simplicity through a Boltzmann function. To achieve adaptation properties, a depleteable vesicle pool model is introduced following Meddis (1986) and Sumner et al. (2002). In this model, the vesicles can exist in three states – free (readily available for release) denoted *q*, released into the synaptic cleft *c* and being reprocessed by the ribbon synapse *w*. Considering *k* being the calcium concentration dependent target release rate, *r* being the re-uptake rate of vesicles from the cleft and *l* the loss rate, *x* the rate of replenishing the free pool and *y* the rate of manufacture, the model can be described mathematically as

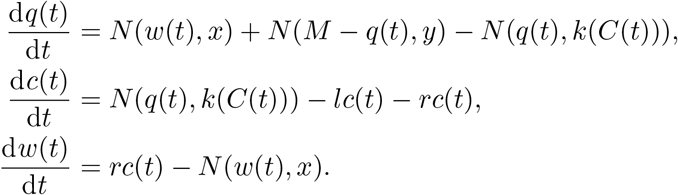

The function *N* (*n, ϱ*) is a stochastic function governing the neurotransmitter transfer of *n* vesicles between the stores, with a mean release rate of *{!* for each single vesicle.

In contrast to the original models by Meddis (1986) and Sumner et al. (2002), our model contains discrete locations for readily releasable vesicles. Therefore, internally, we don’t work with a variable *q* describing a number of readily releasable vesicles, but rather with an array of release sites, for which only valid state transitions may occur (e.g. only an empty release site may be replenished). Following a multivesicular release, the Ca_V_1.3 channel can be temporarily blocked by protons, released from the vesicle to the synaptic cleft (P. F. Vincent et al. 2018). Therefore, a third “blocked” state is added to the two Markov states. However, since we assume the un-blocking of a channel is not affected by external conditions (such as the receptor potential), we can use the Gillespie algorithm, sampling a “blocked duration” randomly upon the blocking of the channel. The probability for the channel to be blocked is assumed to be a Boltzmann function of the normalized proton concentration in the cleft *C*_+_, where *C*_+_ = 1 is the concentration recorded immediately upon a single vesicle release event. The half activation and slope of the Boltzmann function remain calibration parameters, the values 3 and 0.3 yield a reasonable agreement with experiment (P. F. Vincent et al. 2018).

### 5.9 Implementation notes

The model is implemented in MATLAB 2021 optionally employing the Signal Processing toolbox for parts of the analysis and Parallel Computing toolbox for an improved performance. The in-house implementation of the Modified Nodal Analysis (MNA) is based on the symbolic MNA solver SCAM (Cheever 2018). The GNU parallel program (Tange 2018) was used for launching independent simulations in large-scale analysis of the model parameters.

It is worth noting, that our MATLAB implementation allows automatic construction of the final 3D electrical circuit from a single table describing the radial and longitudinal connectivity between electrical elements representing selected cochlear structures. Therefore, the equivalent circuit can be easily modified for studying cochleas of different species or different forms of hearing impairments. An implementation of known deafness mutations (Dror and Avraham 2010; Petit et al. 2001) or changes in aging cochlea such as the reduced number of sensory cells (J. Wang and Puel 2020) are easily implemented.

## Acknowledgements

We would like to thank A. Vetešník, A. Gummer, J. Ashmore, and I. Bruce for sharing implementation of their models which have been used in the development of the present model and served as an important inspiration for us. We are also grateful to S. Johnson for sharing raw data from his IHC patch-clamp measurements, which we used for calibration of the isolated hair-cell model.

**O.T**. acknowledges the Faculty of Mathematics and Physics of the Charles University (Prague, Czech Republic) where he is enrolled as a PhD student and the International Max Planck Research School for “Many-Particle Systems in Structured Environments” (Dresden, Germany) for support. **P.M**. is an employee of Med-El company, which manufactures hearing prosthesis. **P.J**. acknowledges support from the Czech Science Foundation (EXPRO Grant 19-26854X).

## supplementary figures

This document contains supplementary figures for the article “From the ear flap to the nerve: A complete computer model of the peripheral auditory system”. Figure S-1 shows that the model fulfills the zero-crossing condition in response to click stimuli of different sound levels. The figures S-2, S-3, and S-4 contain additional simulation results from the model of the ionic currents within the organ of Corti. The figure S-5 illustrates the difference between classic and stochastic description of mechanosensitive ion channels in the inner hair cells. The figure S-6 shows the effect of voltage dependent gating kinetics of basolateral channels in the IHCs. The figure S-7 verifies, that the high-pass differential equation used for the cochlear amplifier does not affect the results for physiologically relevant frequencies.

**Figure S-1:**
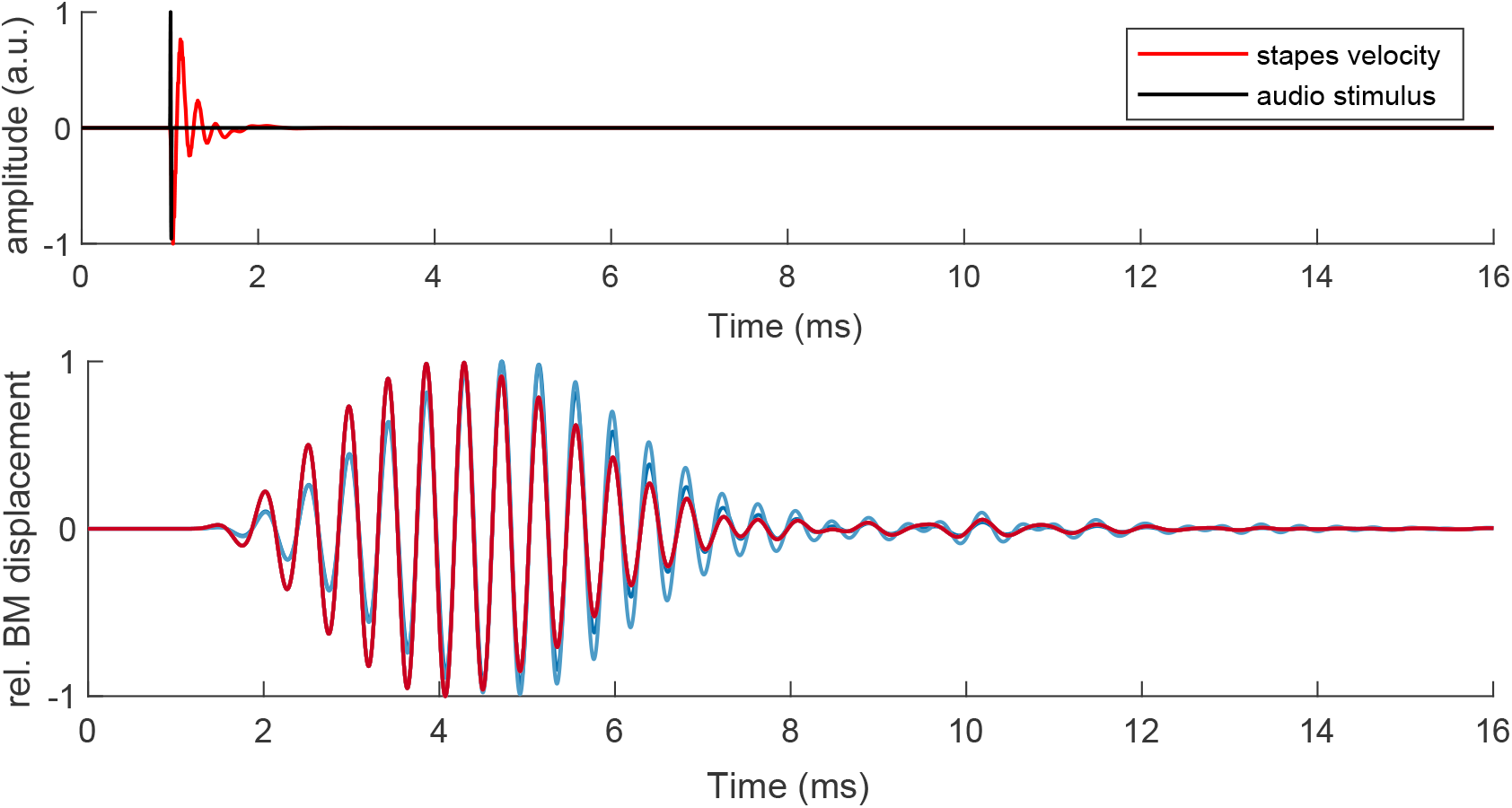
Response of the electro-mechanical model to a click (22 us) of different levels from 0 to 80 dB SPL. **Top:** normalized stimulus waveform with the calculated stapes velocity. **Bottom:** normalized BM displacement at a characteristic position of 1 kHz (as per Li et al. (2021)). While the relative response changes slightly with amplitude, the zero crossing points remain virtually identical, which is consistent with experimental observations.

**Figure S-2:**
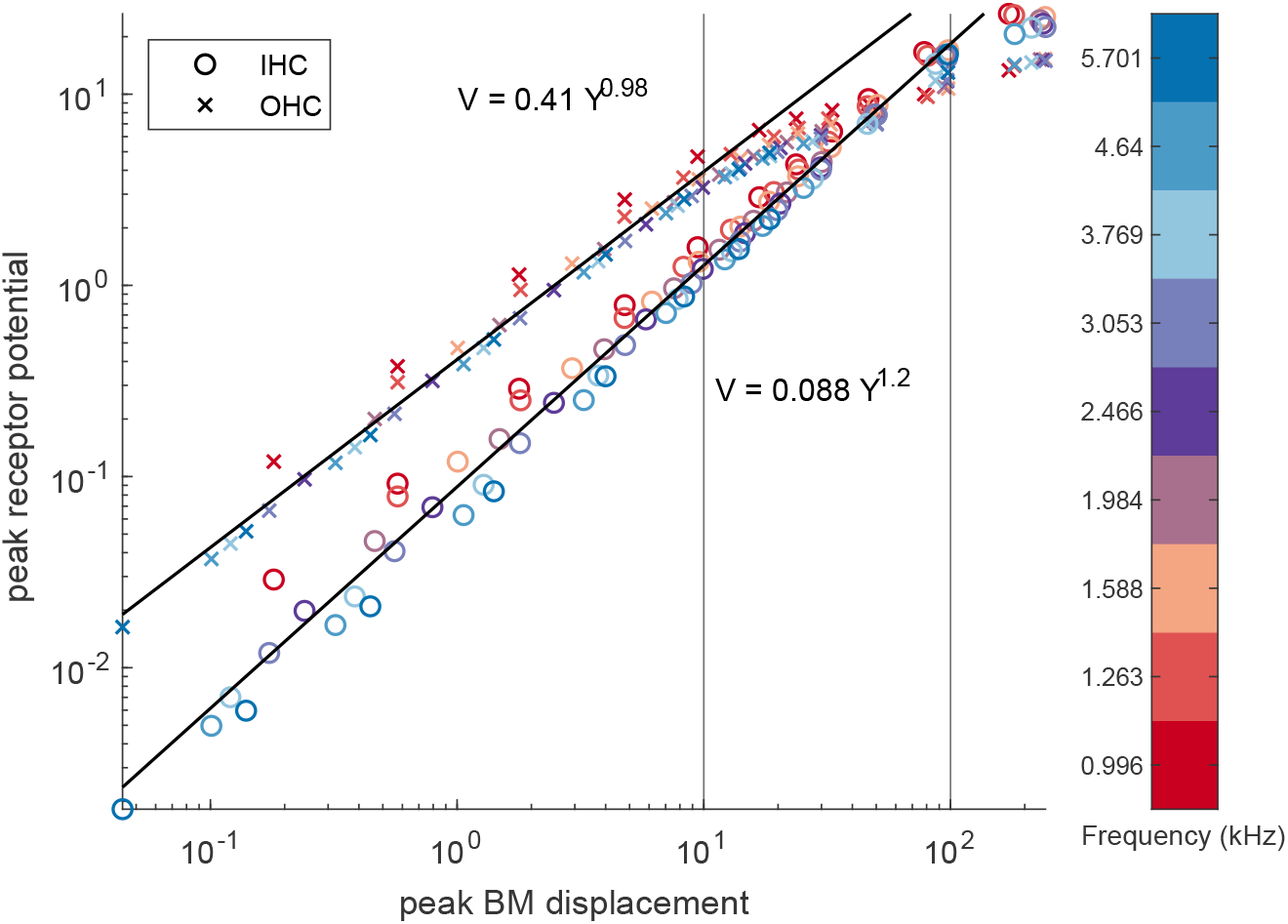
Peak IHC and OHC receptor potential as a function of peak BM displacement (both at the best frequency location) extracted from a set of simulations of pure tones (1–5.7 kHz, 0–100 dB SPL).

**Figure S-3:**
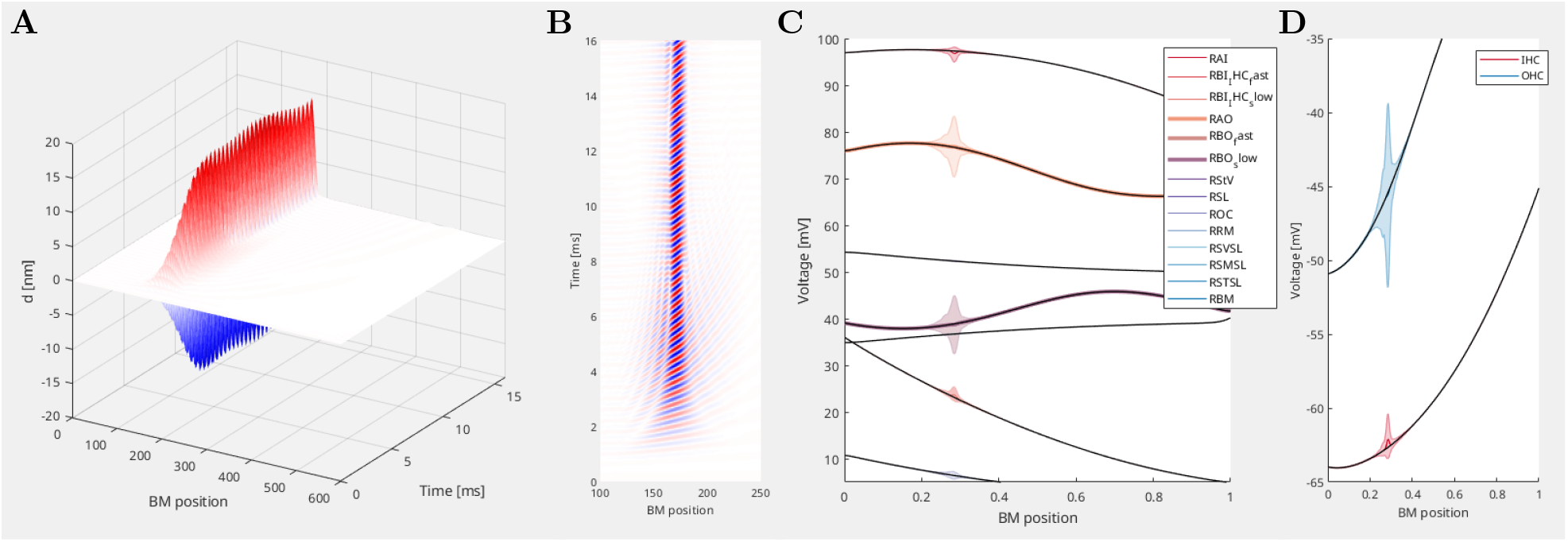
Sample results of the mechanoelectrical model with a pure tone of 2466 Hz and 40 dB SPL. **A-B**: BM displacement as a function of time and position along the BM. The top view (panel B) shows sharpening of the response short after the stimulus onset. **C-D**: Voltage profiles across resistors in the system (C) and voltage across the basolateral IHC and OHC membrane (D). Steady state values in black, changes of sound evoked potentials in colored patches.

**Figure S-4:**
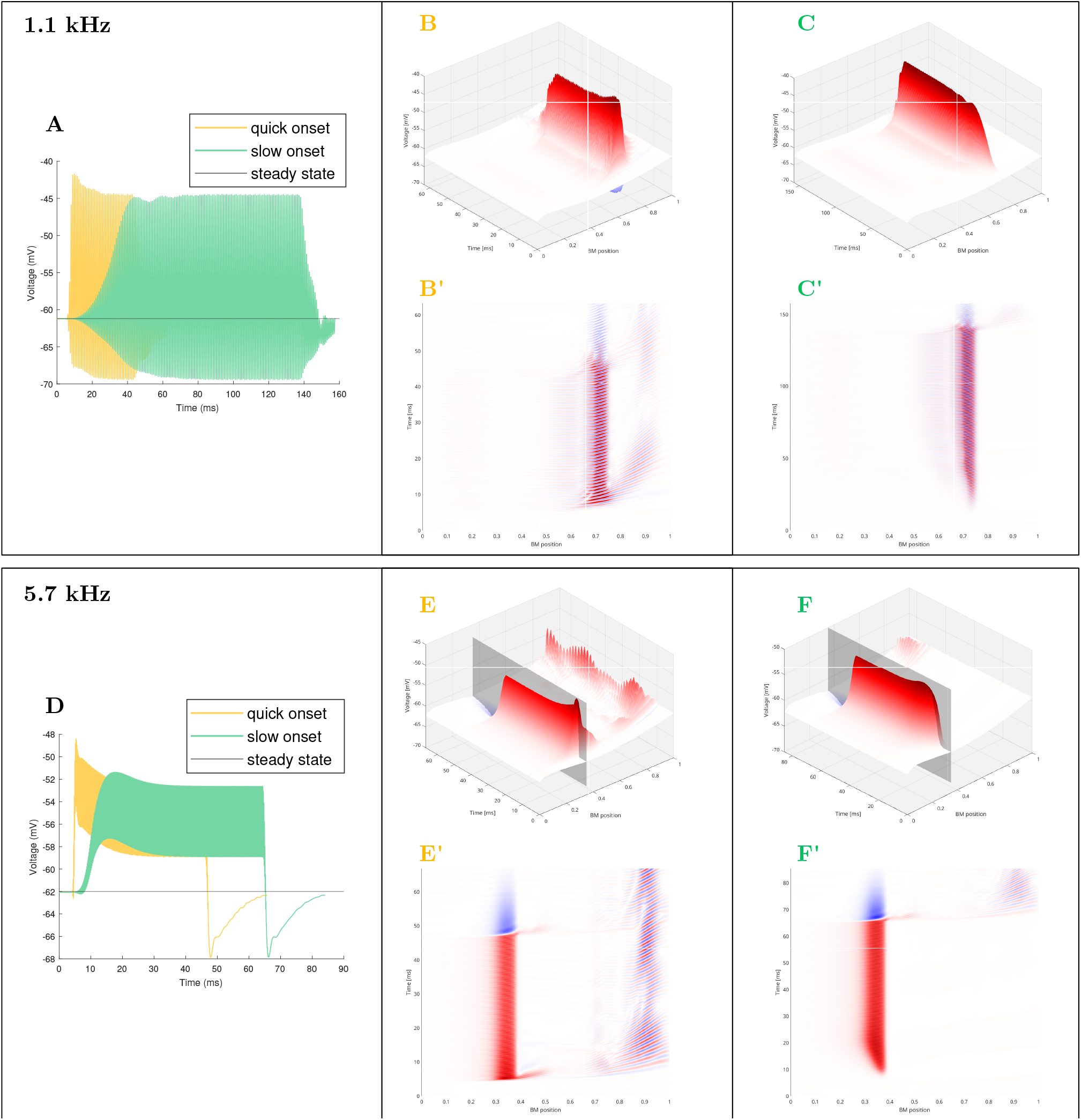
The effect of stimulus onset slope: IHC basolateral transmembrane voltage from four simulations, 5701 and 1123 Hz, 50 dB SPL using a slow and quick onset. **A**,**D**: maximal cross-section, **B**,**C**,**E**,**F**: three-dimensional graph of the voltage, **B’**,**C’**,**E’**,**F’**: “top view”.

**Figure S-5:**
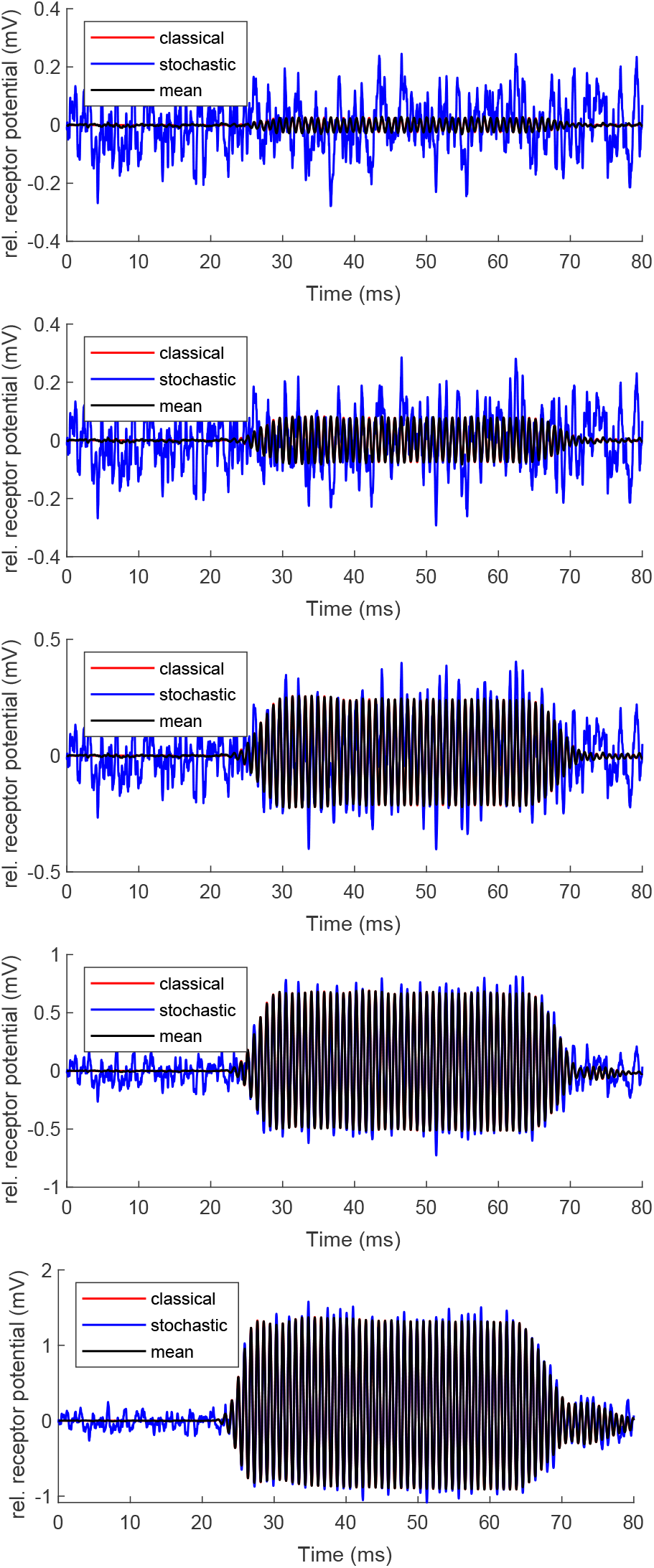
Comparison of the classical and stochastic approach: IHC receptor potential for a 1.1 kHz pure tone of 0, 10, 20, 30, and 40 dB SPL. For low-level stimuli (up to 20-30 dB SPL), the potential of the stochastic model (blue) is significantly different from the continuous model (red) while its mean (1000 independent replicas, plotted black) is almost identical to the continuous model.

**Figure S-6:**
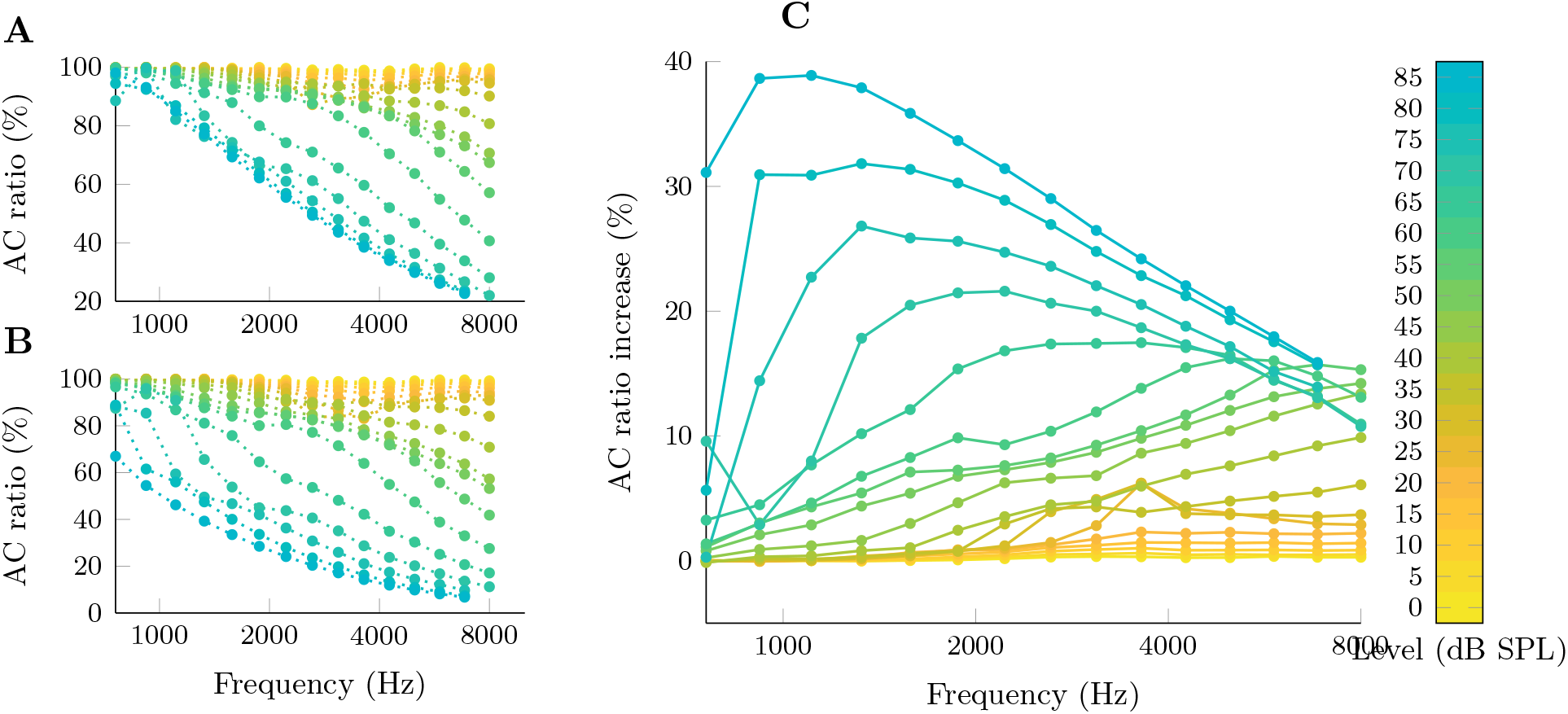
The effect of channel voltage-dependent gating kinetics. IHC receptor potential at the best-frequency location in response to pure tones of varying frequency and level was estimated by the model. The receptor potential can be split into AC and DC component; the AC ratio shown in A–C refers to the magnitude of the AC component relative to the total change of the receptor potential, calculated as AC/(AC + DC). For low level signals the AC ratio (panels A, B) was recorded close to 100 % in the whole frequency range, because the low SPL tones do not elicit significant nonlinearity in the MET channel gating governed by stereocilia deflection. With increasing sound level the MET nonlinearity gains on significance, giving rise to the DC component. For mid- and high-SPL signals, the AC ratio declines faster for high-frequency signals, reaching ∼20 % at 8 kHz and 80 dB SPL. **A**: reference simulation, **B**: simulation in which the IHC basolateral ion channels were considered voltage-independent, **C**: Difference between A and B (calculated as “A” – “B”) showing the increase in the AC ratio that can be attributed to voltage-dependent gating kinetics.

**Figure S-7:**
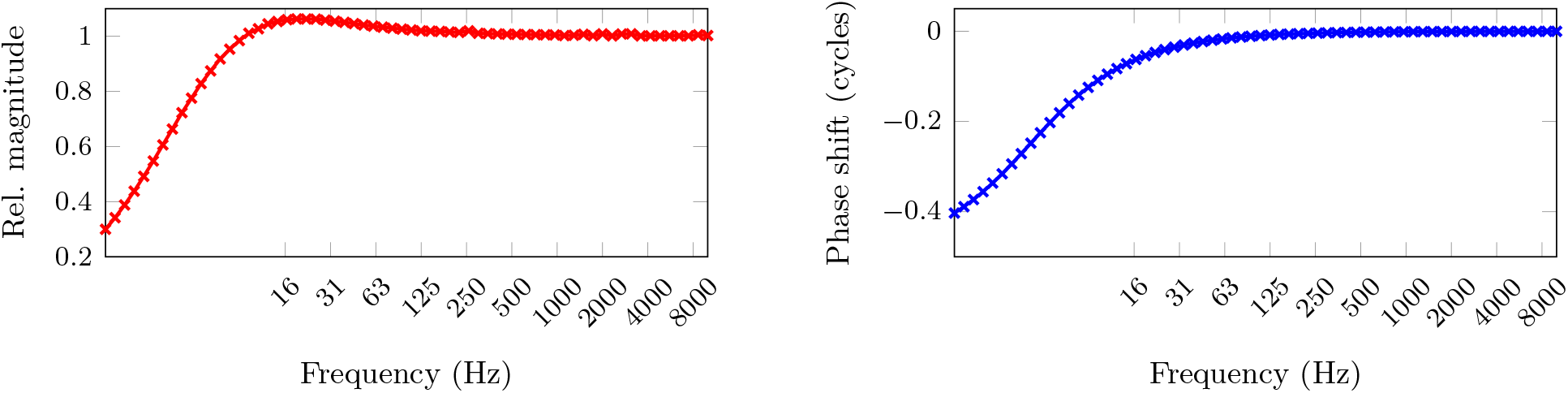
Transfer function of the high pass filtering differential Equation 8. Above 100 Hz, the relative magnitude and phase of the force *F*_ampl_ is the same as the voltage *V*_OHC_.

